# Analysis of chromatin organization and gene expression in T cells identifies functional genes for rheumatoid arthritis

**DOI:** 10.1101/827923

**Authors:** Jing Yang, Amanda McGovern, Paul Martin, Kate Duffus, Xiangyu Ge, Peyman Zarrineh, Andrew P Morris, Antony Adamson, Peter Fraser, Magnus Rattray, Stephen Eyre

## Abstract

Genome-wide association studies have identified genetic variation contributing to complex disease risk. However, assigning causal genes and mechanisms has been more challenging because disease-associated variants are often found in distal regulatory regions with cell-type specific behaviours. Here, we collect ATAC-seq, Hi-C, Capture Hi-C and nuclear RNA-seq data in stimulated CD4+ T-cells over 24 hours, to identify functional enhancers regulating gene expression. We characterise changes in DNA interaction and activity dynamics that correlate with changes gene expression, and find that the strongest correlations are observed within 200 kb of promoters. Using rheumatoid arthritis as an example of T-cell mediated disease, we demonstrate interactions of expression quantitative trait loci with target genes, and confirm assigned genes or show complex interactions for 20% of disease associated loci, including FOXO1, which we confirm using CRISPR/Cas9.

## Introduction

It is now well established that the vast majority of SNPs implicated in common complex diseases from genome-wide association studies (GWAS) are found outside protein coding exons and are enriched in both cell type and stimulatory dependent regulatory regions^1,2^. The task of assigning these regulatory enhancers to their target genes is non-trivial. First, they can act over long distances, often ‘skipping’ genes^3^. Second, they can behave differently dependent on cellular context^4,5^, including chronicity of stimulation^6^. To translate GWAS findings in complex disease genetics, one of the pivotal tasks is therefore to link the genetic changes that are associated with disease risk to genes, cell types and direction of effect.

Popular methods to link these ‘disease enhancers’ to genes is either to determine physical interactions, with methods such as Hi-C^7^, use quantitative trait analysis^4^ or examine correlated states^8^, with techniques such as ChIP-seq and ATAC-seq, linked to gene expression. The vast majority of these studies, to date, have investigated these epigenomic profiles at either discrete time points^9,10^ (e.g. baseline and/or after stimulation), and/or by combining data from different experiments (e.g. ATAC-seq and Hi-C)^9^.

Over 100 genetic loci have been associated with rheumatoid arthritis (RA), a T-cell mediated autoimmune disease. Of these, 14 loci have associated variants that are protein-coding and 13 have robust evidence through expression quantitative trait locus (eQTL) studies to implicate the target gene. The remainder are thought to map to regulatory regions, with so far unconfirmed gene targets, although we, and others, have previously shown interactions with disease implicated enhancers and putative causal genes^3,11,12^.

Here we have combined simultaneously measured ATAC-seq, Hi-C, Capture Hi-C (CHi-C) and nuclear RNA-seq data in stimulated primary CD4+ T cells (Fig. 1), to define the complex relationship between DNA activity, interactions and gene expression. We then go on to incorporate fine-mapped associated variants from RA, and validate long range interactions with CRISPR/Cas9, to assign SNPs, genes and direction of effect to GWASloci for this T-cell driven disease.

**Fig. 1.**
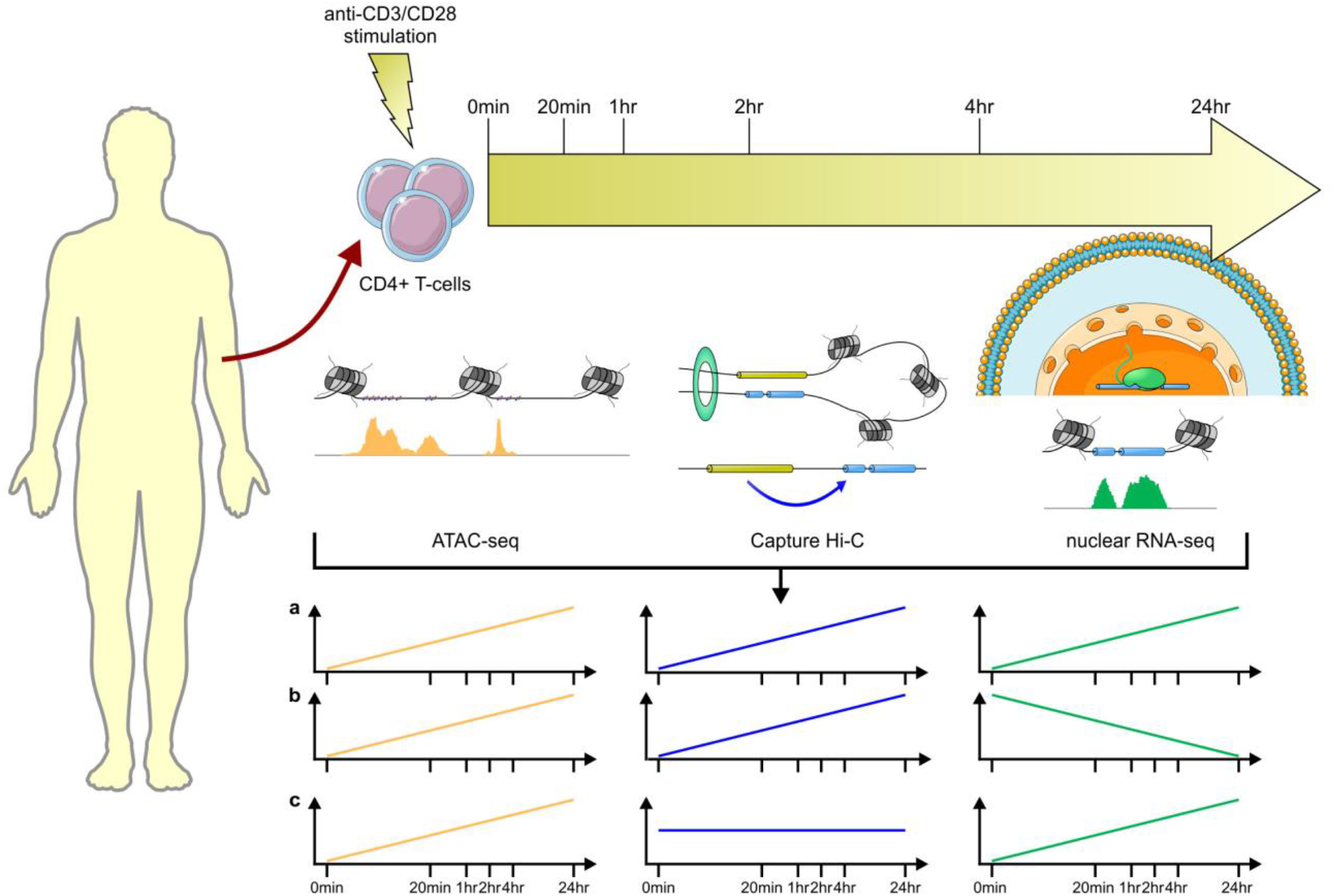
Schematic of the study design. ATAC-seq, CHi-C and nuclear RNA-seq experiments were carried out for unstimulated and stimulated CD4+ T-cell samples at time 0 mins, 20 mins, 1 hr, 2 hrs, 4 hrs and 24 hrs. Time course profiles were created by aligning features (ATAC-seq peaks and CHi-C interactions) across time and counting reads supporting each feature at each time point.

## Results

### High-quality data from sequenced libraries

A total of 116.7 ± 28.5 million reads per sample were mapped for RNA-seq by STAR^13^ with alignment rates over 98% across six time points: 0 mins, 20 mins, 1 hr, 2 hrs, 4 hrs and 24 hrs after stimulation with anti-CD3/anti-CD28. A total of 76,359 ATAC-seq peaks were obtained; 6,287 peaks unique to unstimulated cells; 45,869 only appearing after stimulation; and 24,203 shared across all time points. A total of 271,398 CHi-C interactions were generated from the time course data and interactions were retained as features when at least one time point showed a significant interaction. Of these interactions, 94% occurred within the same chromosome and 57% were within 5 Mb of promoters.

Gene expression data demonstrated good correlation between replicates (Supplementary Fig. 1a-d). From an initial FACS analysis (Supplementary Table 1), and comparison of RNA-seq data, we determined the initial T cell purity of the samples to be around 94% CD4+, displaying similar amounts of CD4+ memory and CD4+ naïve (Supplementary Fig. 2a). To assess the amount of stimulation we explored markers of naivety, activation and cytokine expression and confirmed a similar degree of stimulation across all the samples (Supplementary Fig. 2b).

ATAC-seq peaks also demonstrated good quality and correlation between replicates (Supplementary Fig. 3a-e) and enrichment for both marks of enhancer activity (ChromHMM, Supplementary Fig. 4a-d) and CTCF sites across all time points (Supplementary Fig. 4e), with an increased amount of CTCF occupancy in ATAC-seq peaks at TAD boundaries, as expected (Supplementary Fig. 4f).

### Data consistency with previous studies

Comparison of ATAC-seq peaks from CD4+ baseline (unstimulated) and 48 hrs after stimulation peaks from a similar dataset^9^ revealed strong concordance (Supplementary Fig. 3f): 71% of peaks (21,549/30,403) from unstimulated CD4+ T cells overlapped in both datasets^9^, while 75% of peaks (22,911/30,593) from stimulated CD4+ T cells (24 hrs vs 48 hours post stimulation) overlapped ^9^. We also observed a similar magnitude of increase in the number of ATAC-seq peaks for merged unstimulated and stimulated data.

Comparison of our data with published datasets from the same cell-type and stimulation revealed CHi-C data, both unstimulated or stimulated for 4 hrs, demonstrated good consistency (Supplementary Fig. 5a). When restricting all the interactions from both unstimulated and stimulated to those that share the same baits, we found 57% of interactions (27,794/48,570) to overlap (by at least 1 bp) between our study and previously identified interactions^10^ and this increases to 73% for interactions within 5 Mb of promoters (26,836/36,698) and 87% within 200 kb of promoters (8,367/9,627). This strongly suggests that the interactions between promoters and active enhancers within 200kb are consistent, robust and reproducible between studies. We found 22,126 genes with evidence of CD4+ T cell expression for at least one time point in our RNA-seq data. We considered genes classified as “Persistent repressed”, “Early induced”, “Intermediate induced I”, “Intermediate induced II” and “Late induced” in a previous study^4^, and found that these genes exhibited similar patterns of expression in our RNA-seq data (Supplementary Fig. 1e-i).

### Chromatin conformation dynamics

It is well established that gene expression changes with time after stimulation in CD4+ cells^4^ and we find similar changes to previous studies, with a range of dynamic expression profiles corresponding to genes activated early, intermediate or late, or repressed (Supplementary Fig. 1e-i). However, it is less well established how chromatin structure changes post stimulation, in the form of A/B compartments, topologically associating domains (TADs) and individual interactions, or how enhancer activity and open chromatin change over time.

Based on Hi-C matrices with resolutions of 40 kb, 1,230 merged TADs were recovered across all the time points with an average size of 983.2 kb. Consistency of TADS across time points was measured by the fraction of overlapping TADS (90% reciprocal), over the number of TADs at each specified time point (Supplementary Fig. 6a). On average between time point 0 and 4 hours 84% of TADS intersect. When adding the 24 hour TAD data, the mean intersection reduces to 74%, illustrating more substantial dynamic changes in TADs over longer times. Fig. 2b shows the stratum adjusted correlation coefficient (SCC) between Hi-C datasets^14^ and shows a slight but significant reduction in correlation as the time separation of experiments increases, consistent with our observations regarding TADs. SCC between replicates and within replicates does not show clear differences, implying that the changes in correlations are not due to batch effects.

**Fig. 2.**
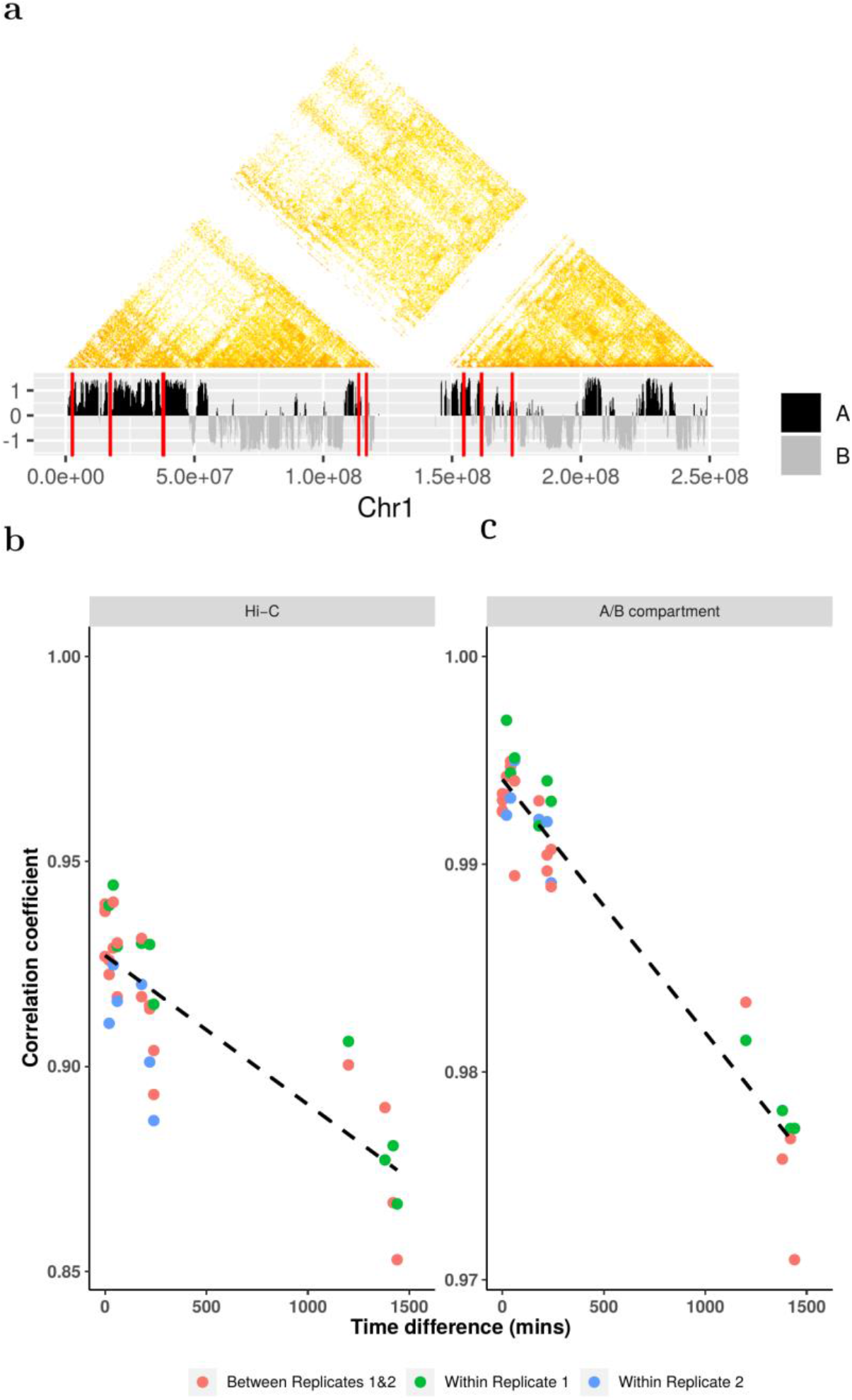
Illustration of Hi-C dynamics. **a**, Hi-C interaction matrix (40 kb resolution) of replicate 1 for chr1 at time 0 mins (upper) with corresponding A/B compartments (lower), where red lines represent positons of SNPs. **b**, Correlation changes between Hi-C data with respect to differences between times, where the dashed line is the fitted linear line for the correlation coefficients. **c**, Correlation changes of A/B compartments with respect to the differences between times, where the blue line shares the same information as conveyed in the plot b.

We recovered 1,136 A compartments and 1,266 B compartments merged across the time course data, with the maximum compartment sizes being 39.5 Mb and 34.5 Mb, respectively. Similar to TADs, the consistency of compartments across time points was measured by the fraction of overlapping compartments (90% reciprocal) over the number of compartments at each specified time point (Supplementary Fig. 6 b,c). The same consistency was observed for compartments A and B. Fig. 2c shows the correlation between A/B compartment allocations, demonstrating a slight but significant reduction between experiments as time separation increases. Compartments that changed over time were found to be enriched for lamina associated domains (LADs) and genes that were ‘intermediate’ or ‘late’ expressed (Supplementary Fig. 7).

These results are broadly consistent with other studies, demonstrating how the higher chromatin conformation states, in the form of A/B compartments and TADS, is largely invariant between cell types^11^. Here, we demonstrate similar levels of consistency in a single cell-type post stimulation (Supplementary Fig. 8a-c).

In contrast to the relative invariance of TADs and A/B compartments, our CHi-C data, analysing interactions between individual restriction fragments, showed a much greater degree of dynamics. We used the Bayesian Information Criterion (BIC)^15^ and a χ² test to compare a dynamic (Gaussian process) model to a static model^16^ for CHi-C interaction count data across time. We found 24% (63,843/271,398) of CHi-C links with evidence of change over time (BIC_dynamic_ < BIC_static_) and 7.5% of interactions showed stronger evidence of change over time (20,224/271,398, χ² test, *p* < 0.05), among which 24% (4,837/20,224) are within 200 kb of promoters.

### Open chromatin dynamics

We compared a dynamic (Gaussian process) and static model for ATAC-seq time course data to identify changes in open chromatin across time and found 11% (7,852/74,583) of ATAC-seq peaks with evidence of change over 24 hrs (BIC_dynamic_ < BIC_static_) with 2,780 of these peaks showing stronger evidence of change (χ^2^ test, *p* < 0.05). A heatmap of ATAC-seq time course data (Fig. 3a) demonstrates six broad patterns of change (Fig.3b). Mapping of transcription factor binding sites (TFBS) motifs under these broad clusters revealed a strong enrichment of transcription factors known to be important in CD4+ stimulation and differentiation. The AP-1 TFBS (e.g. BATF) motif was shown to be enriched in low to high activity, whilst strong enrichment of ETS/RUNX1 TFBS was seen in models of high to low activity, and a strong enrichment of CTCF and BORIS motifs was observed in the models that demonstrated transient dynamics before returning to baseline after 24 hours. These findings match those reported in a previous study of ATAC-seq data in CD4+ T cells stimulated with anti-CD3/anti-CD28^9^. There it was demonstrated that the AP-1/BATF motif was enriched in stimulated ATAC-seq peaks, ETS/RUNX in unstimulated cells and CTCF/BORIS motifs were detected under the ‘shared’ unstimulated and stimulated peaks, closely matching our findings. ATAC-seq peaks overlapping H3K27ac marks and without CTCF binding are more likely to be gained over time (blue to red, Supplementary Fig. 9a), a pattern not seen in ATAC-seq peaks bound by CTCF without H3K27ac, although the distribution of interactions from these two classes of peaks (gained or lost) remains similar, with no bias seen in either category of peak (Supplementary Fig. 9b). We found evidence that the increase in ATAC-seq activity over time (e.g. Fig. 3b, cluster 1), corresponded to an increase in interactions (Supplementary Fig. 9d), particularly DNA locations containing the JunB transcription factor.

**Fig. 3.**
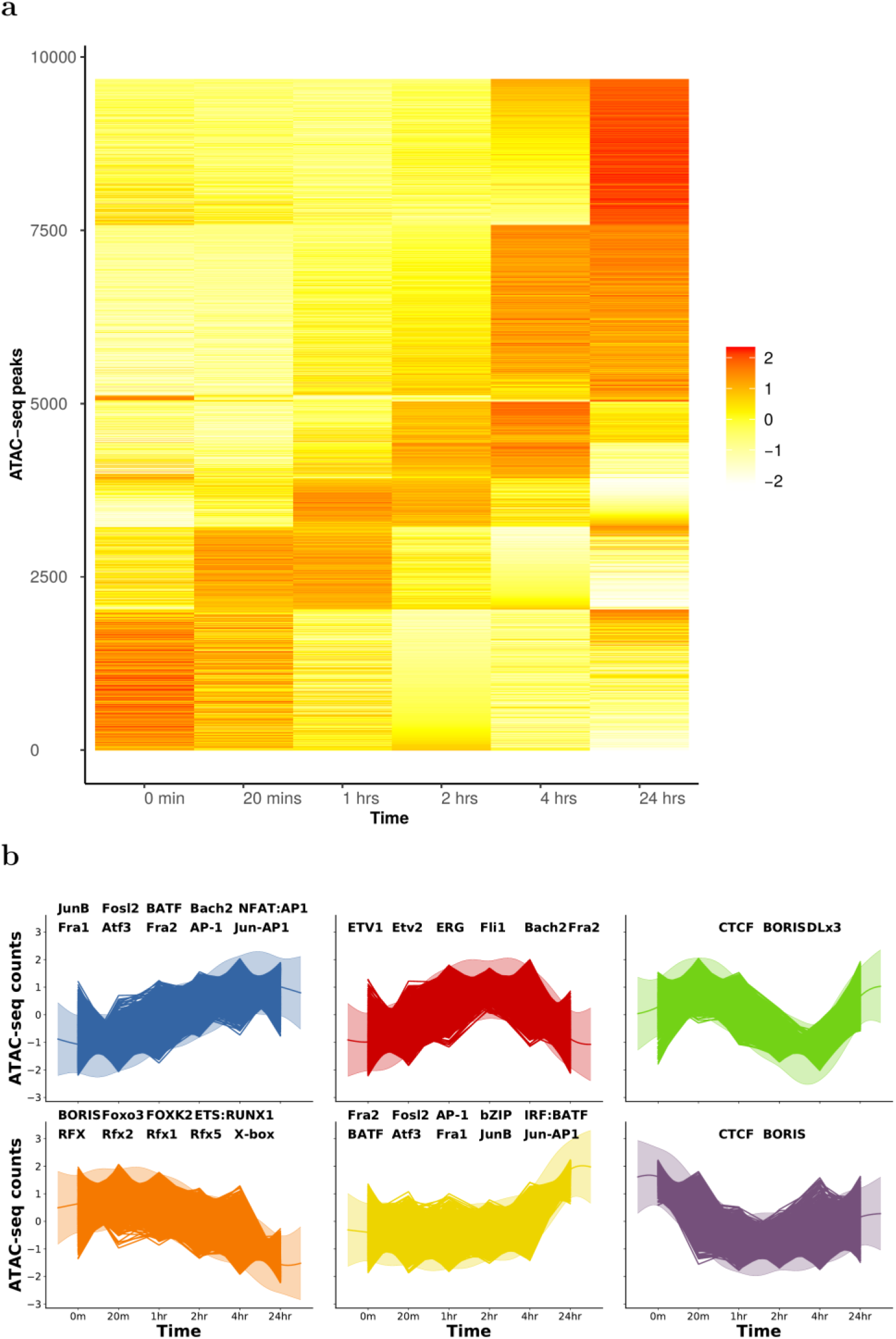
Illustration of ATAC-seq time course profile dynamics. **a**, heatmap of ATAC-seq counts data for peaks showing evidence of temporal dynamics. **b**, Clustering of ATAC-seq time course data using a Gaussian process mixture model. Significantly enriched DNA-binding MOTIFs in each peak (using static peaks as background) are labelled in each cluster.

### Correlating chromatin dynamics with gene expression

We next went on to test whether these dynamic measures of DNA activity, interaction and expression exhibited any correlation between their time course profiles. Previous studies, using measurements of H3K27ac, Hi-C and expression across different cell types, demonstrated how subtle changes in contact frequency correlated with larger changes in active DNA and expression^17^. We wanted to determine the nature of this relationship in our data, from a single, activated cell type. We looked for otherEnd baits that contained ATAC-seq peaks, and linked these ATAC-seq peaks with CHi-C interactions to promoter baits and associated genes. Using this approach, we formed 37,819 links. Pearson correlation coefficients for paired time course data between ATAC-seq and CHi-C, ATAC-seq and gene, and gene and CHi-C were calculated. We used a link randomisation procedure to identify whether the number of correlations observed at a particular level could be considered significantly enriched (see Methods). We show an enrichment for extreme correlations between ATAC-seq, CHi-C and RNA-seq datasets, particularly an enrichment for high positive correlations within 200 kb of promoters (Fig. 4a), suggesting that functional, interactive correlations are most common within ‘contact domains’. This observation is supported by previous findings, where the median distance between H3K27ac loop anchors and interacting otherEnds (130 kb)^5^ and the median distance of cohesion constrained regulatory DNA-loops (185 kb)^7^ are typically within a ~200 kb range. Boxplots of the log fold change in ATAC-seq, CHi-C and RNA-seq intensity in the highly correlated regions (Fig. 4b,c) revealed how relatively small changes in both ATAC-seq and CHi-C intensity (~2 fold change) correlated with larger changes in expression (~2 fold change). This is consistent with similar patterns observed in different cell types^17^.Previous studies have indicated how i) using eQTL data, ~50% of ATAC-seq peaks are already active/poised before influencing gene expression^18^, ii) using HiChIP data, expression can be correlated with either H3K27ac or interactions^5^, and iii) empirical ranking of enhancers by CRISPR corresponds most strongly when combining terms for interaction and activity^19^,. Taken together, these observations suggest that both interactions and activity have important roles in gene regulation. Examining the relationship of CHi-C interactions, DNA dynamics and expression in closer detail in our data with 200 kb distance between bait and otherEnd fragments revealed three broad patterns of dynamics associated with four clustered gene expression patterns (Supplementary Fig. 10): Approximately 8% (469/5,939) of links were associated with dynamic ATAC-seq peaks only (Supplementary Fig. 10a,b), 32% (1,901/5,939) were associated with dynamic CHi-C interactions only (Supplementary Fig. 10c,d) and 6% (349/5,939) were associated with dynamics in both (Supplementary Fig. 10e,f). Our findings, together with previous studies, therefore suggest that both activity and interactions are independently important in gene regulation and that subtle change in interaction and ATAC-seq intensity have a larger effect on gene expression.

**Fig 4.**
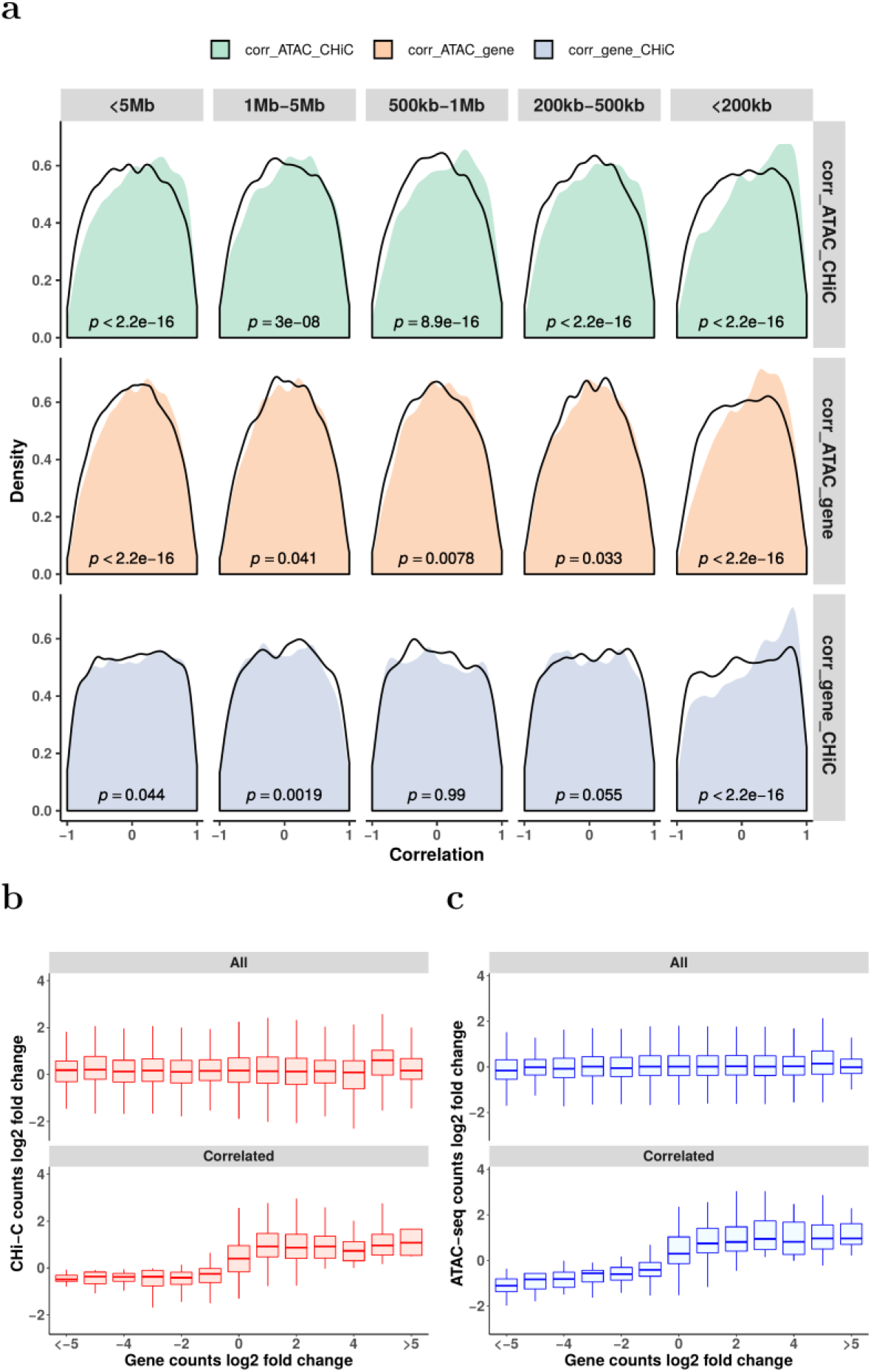
Illustrations of the correlations between ATAC-seq, CHi-C and RNA-seq time course profiles. **a**, Density plots of the Pearson correlation coefficients between ATAC-seq and CHi-C, ATAC-seq and gene and gene and CHi-C under various distance ranges around promoters. Distances ranges include those less than 5 Mb, between 1 Mb and 5 Mb, between 500 kb and 1 Mb, between 200 kb and 500 kb and less than 200 kb. Black lines represent the density plots of the corresponding random background. P values from Wilcoxon tests are labelled in each panel. **b**, Comparison of the log2 fold change between CHi-C and gene data for all dataset (upper) and those highly correlated ones with Pearson correlation coefficients between ATAC-seq, gene and CHi-C over 0.5 (lower). **c**, Comparison of the log2 fold change between ATAC-seq and gene data for all datasets (upper) and those highly correlated ones with Pearson correlation coefficients between ATAC-seq, gene and CHi-C over 0.5 (lower).

### Prioritisation of causal genes in GWAS loci for autoimmune disease

We identified 312 loci which contained an autoimmune disease lead SNP that was highly correlated with a variant (r^2^>0.8) located within an ATAC-seq peak, which was itself interacting and highly correlated with the expression of a gene (Supplementary Table 3). These data highlight potential causal SNP, ATAC-seq and gene relationships for autoimmune disease risk in CD4+ T-cells, confirming previous reports or providing evidence for genes such as *PRKCQ, CD44, ETS1* and *ARID5B*.

We next considered 80 of the 100 loci previously associated with RA that attain genome-wide significance in European ancestry GWAS meta-analysis^20,21^. For each locus, we constructed a 99% credible set of SNPs that accounted for 99% of the probability of containing the causal variant. We found that 97% (2,131/2,192) of RA-associated variants from our 99% credible SNP sets lie within A compartments across all time points while only 28 (1%) lie consistently within B compartments after stimulation. Fifteen credible set SNPs were found in regions that change between A and B over time. These included RA-credible set SNPs on chromosome 1, proximal to the *TNFSF4* and *TNFSF18* genes, that were initially contained within an inactive B compartment at 20 mins and were then found in an A compartment at 4 hrs (Supplementary Fig. 8d).

We next investigated whether we could map RA GWAS implicated ATAC-seq peaks to genes, identify the likely causal SNPs within the peaks and determine a mechanism and direction of effect through the correlated expression data. Genome-wide, RA credible set SNPs signals were enriched (5-30 fold) in open regions of chromatin, as expected. The strongest enrichment was observed at 4 and 24 hours post stimulation (Supplementary Fig. 11a), where SNPs mapping to open regions of chromatin explained up to 30% of the heritability of RA (Supplementary Fig. 11b). We found that 43/80 GWAS loci contained 67 ATAC-seq peaks with at least one RA credible set SNP SNP (96 SNPs in total) that interacted and correlated with the expression of 168 genes, an average of 2.6 genes per peak (Supplementary Fig. 12 and Supplementary Table 3). These data have the ability to limit the number of putative causal SNPs/ATAC-seq peaks for each locus. The 43 loci with ATAC-seq peaks that contain RA GWAS associated SNPs have 5527 credible SNPs, which reduce down to 98 SNPs in 67 ATAC-seq peaks that interact with and are correlated with the expression of 168 genes. For example, there are 18 SNPs within the 99% credible SNP set for the *RBPJ* locus, but this reduces to 2 SNPs within 2 ATAC-seq peaks that interact and correlate with gene expression (Supplementary Fig 13a). For 6 loci interactions between 30-220kb implicate either a single ATAC-seq peak (*CD5, PXK, TCTE1, CDK6, TPD52*) or 2 ATAC-seq peaks (*IL6ST*) that contain an RA credible set SNP, interacting with the target eQTL gene. This implicated a single likely causal SNP in 3 loci (*PXK, CDK6, TPD52, CD5*) and less than 3 likely causal SNPs in the other loci (*IL6ST, TCTE1*) (Supplementary Table 3).For 13 loci, our correlated, dynamic data support the currently assigned gene, and provide the first biological evidence for 7 of these genes (Supplementary Table 3). These genes include *DDX6, PRKCH, RBPJ, PVT1, PRDM1, and PTPN2*, with many interactions confirming genes up to 200 kb, and one over 800 kb (*PVT1*), from the RA credible set SNPs SNP, but always constrained within TADs.

For a number of loci, our results point to complex relationships between RA credible set SNP regions and putative causal genes (Supplementary Table 3). For some regions, whilst we did not demonstrate direct interaction between an ATAC-seq peak and gene, the genomic region spanning the credible SNP set made a long distance physical interaction with genes that have supportive evidence for disease involvement. For example, on chromosome 10, an intronic region within the *ARID5B* gene, containing RA credible set SNPs, interacts with *RTKN2,* involved in the NFKB pathway, and containing nsSNPs associated with Asian RA^22^ (Supplementary Fig 13b). Similarly on chromosome 3, a region with RA credible set SNPs intergenic of *EOMES* interacts with *AZI2* (an activator of NFKB), suggesting the region contains an enhancer that could potentially control two important genes in the T cell immune pathway (Supplementary Fig. 13c). We also provide evidence for complex regions with direct ATAC-seq, gene promoter interactions. Two regions in particular provided insight into potential genes for RA. Two ATAC-seq peaks, containing 5 SNPs that are RA credible set SNPs on chromosome 8, both interact with *PVT1* and *MYC1*, situated some 450-800 kb away from these peaks (Fig. 5). Here we demonstrate a positive correlation between the ATAC-seq peak dynamics and gene expression for both *PVT1* and *MYC1* (r^2^=0.67 and 0.68 respectively). Interestingly, this ATAC-seq peak region has previously been demonstrated to be a key repressor of *MYC1* gene expression, following a comprehensive CRISPRi screen in K562 erythroleukemia cells^19^. We therefore confirm this relationship between a distant regulatory region and *MYC1* expression in primary T cells, highlighting the likelihood that this gene has a role in the susceptibility to RA.

**Fig. 5.**
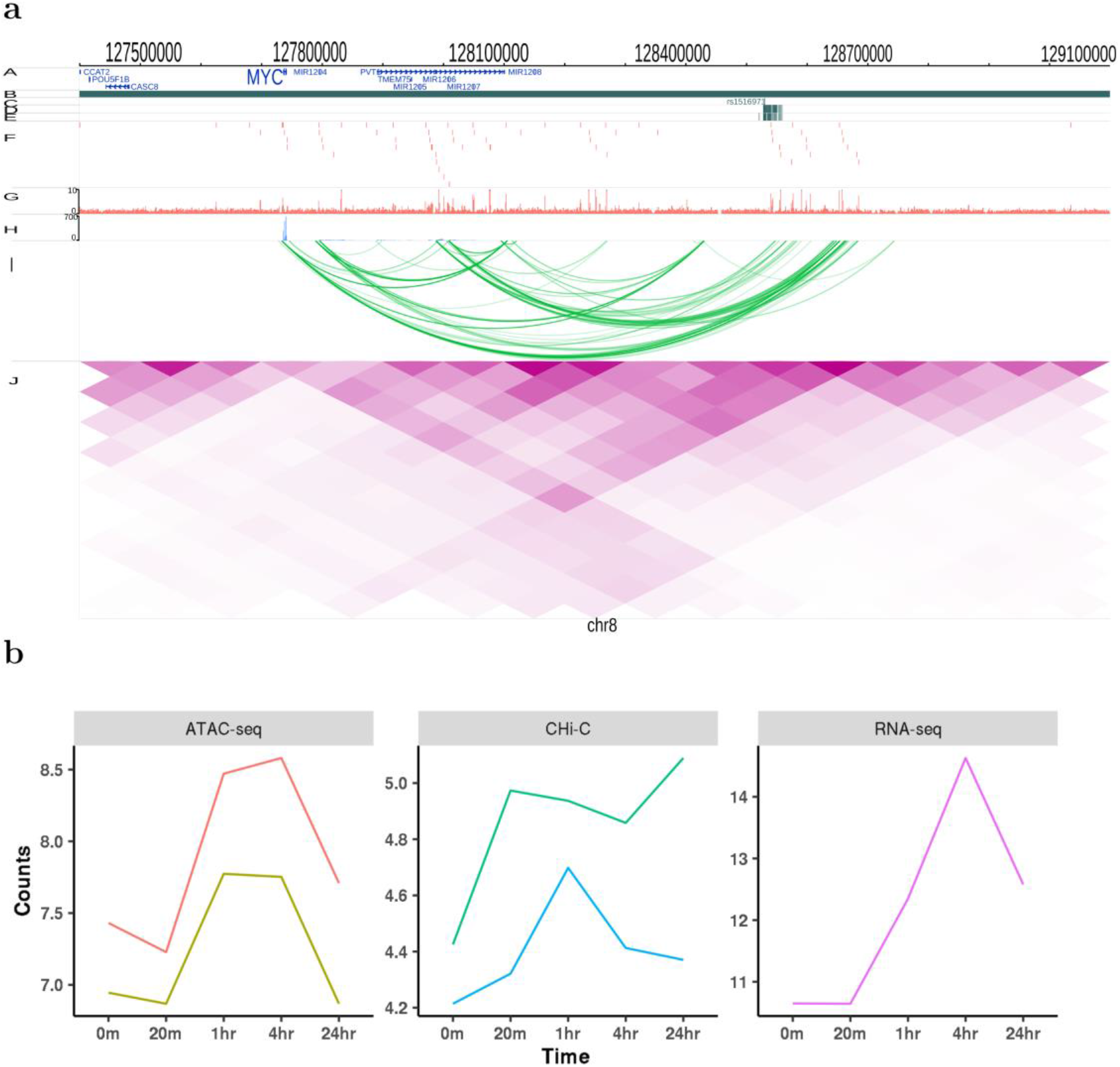
Illustration of genomic interaction activities around MYC. **a**, Screenshot of the SNPs (dark green), ATAC-seq peaks (red), RNA-seq (lightblue) and CHi-C interactions (green) and heatmap from Hi-C (purple) around MYC at time 4 hrs. yaxis labels A to J represent different tracks plotted. A: RefSeq genes, B: TADs, C: Index SNPs, D: LD SNPs, E: 99% credible sets, F: ATAC-seq peaks, G: ATAC-seq signal, H: RNA-seq, I: CHi-C, J: Hi-C. **b**, Time course profiles of ATAC-seq (left), CHi-C (middle) and RNA-seq (right) of the data associated with SNPs data around MYC.

Putting this gene list through a drug target pipeline^23^ demonstrated how the newly implicated or confirmed genes from this study, such as *MyC*, *GSN* and *MMP9* are existing therapeutic targets (Supplementary Table 2)

### Confirmation of correlated interaction with CRISPR/Cas9

Finally we sought to demonstrate how an ATAC-seq peak, containing anRA credible set SNP, intronic of the *COG6* gene interacts with, and is correlated with the expression of, the *FOXO1* gene located some 900 kb away (Fig 6). We investigated whether this dynamic ATAC-seq peak is functionally interacting with the *FOXO1* promoter, a transcription factor involved in T cell development, and a gene that has previously been strongly implicated in RA through functional immune studies in patient samples^24–27^. To do this we used CRISPRa, with dCas9-p300, and the HEK cell line. Although using a cell line, as opposed to primary immune cells, is not optimal, we used HEK cells as the model system since we and others have previously demonstrated how the TAD region containing both *COG6* and *FOX01* is highly conserved in a wide range of cell types, including primary cells as well as various cell lines (Jurkat, GM and HEK). We designed guide RNA to cover the single enhancer that contains the 2 ATAC-seq peaks and 3 RA credible set SNPs, pooling the guides in a single transfection (Supplementary Fig. 14). We demonstrated that, when we activate the *COG6* intronic enhancer with this system and targeted gRNAs, not only do we observe a consistent increase in *COG6* mRNA expression itself, we obtain robust, reproducible up regulation of *FOXO1* gene expression (Fig. 7). Although the credible set SNP in this region is a strong eQTL for *COG6*, this CRISPR validation of the correlated interaction, DNA activity and expression data, alongside previous immunological studies, implies that the associated enhancer may have diverse roles on a number of genes within this 1 Mb TAD region, and that GWAS implicated enhancers should not necessarily be assigned to single genes.

**Fig. 6.**
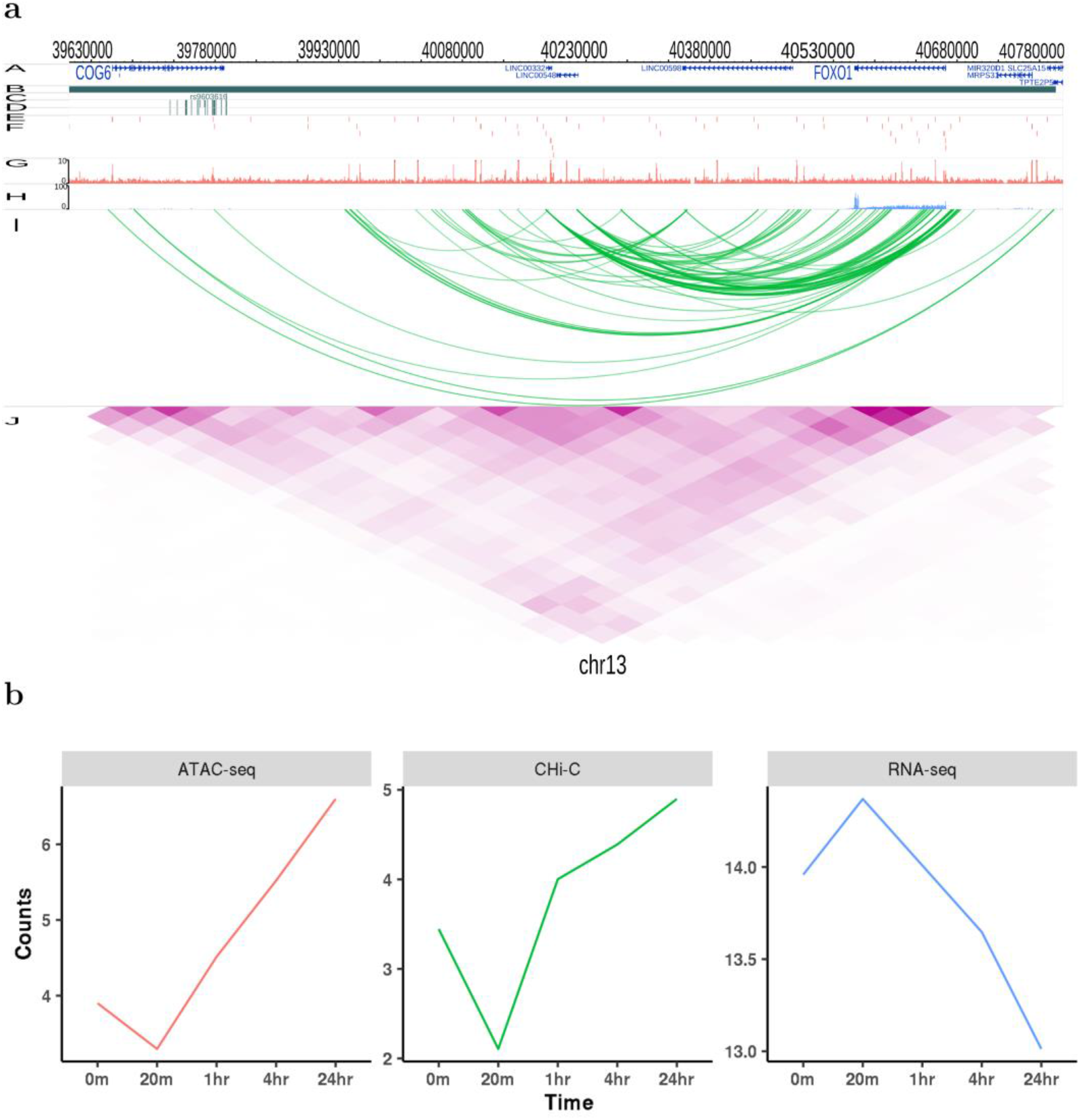
Illustration of genomic interaction activities around FOXO1. **a**, Screenshot of the SNPs (dark green), ATAC-seq peaks (red), RNA-seq (lightblue) and CHi-C interactions (green) and heatmap from Hi-C (purple) around FOXO1 at time 4 hrs. yaxis labels A to J represent different tracks plotted. A: RefSeq genes, B: TADs, C: Index SNPs, D: LD SNPs, E: 99% credible sets, F: ATAC-seq peaks, G: ATAC-seq signal, H: RNA-seq, I: CHi-C, J: Hi-C. **b**, Time course profiles of ATAC-seq (left), Chi-C (middle) and RNA-seq (right) of the data associated with SNPs data around FOXO1.

**Fig. 7.**
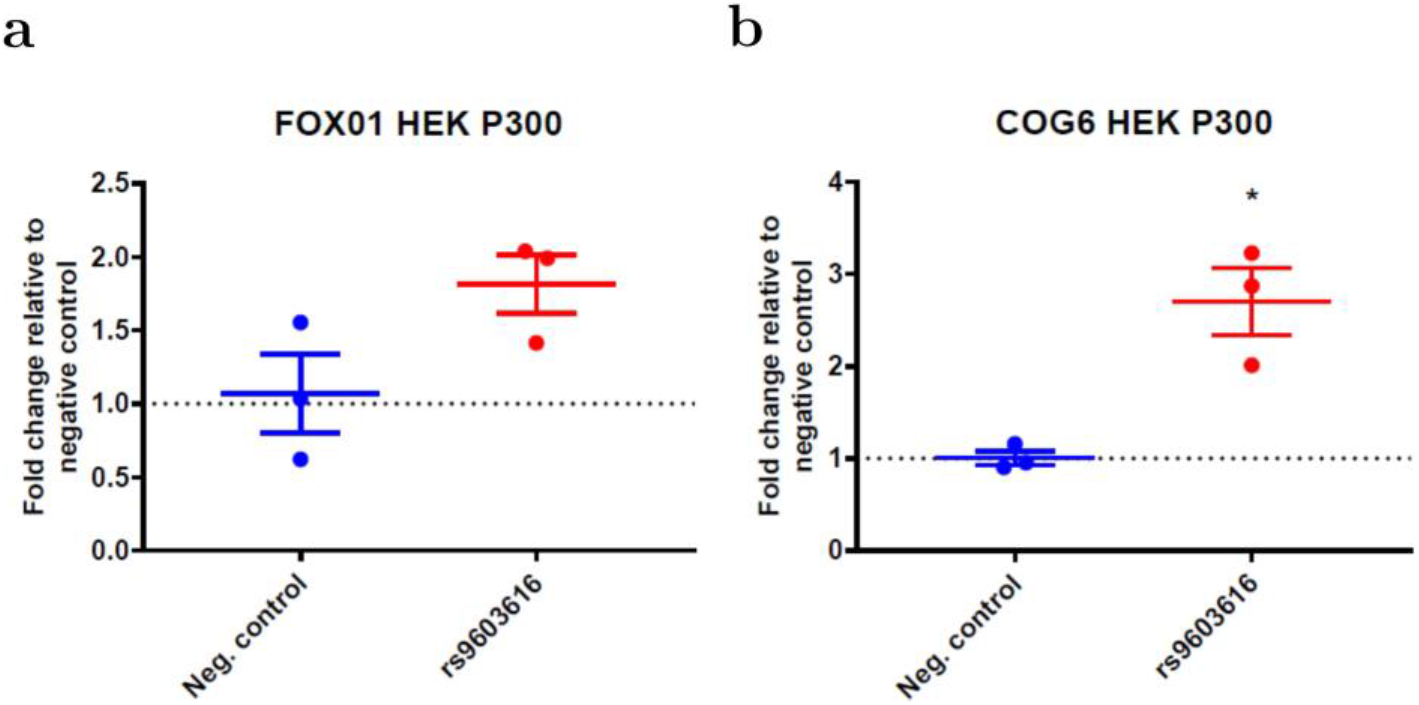
CRISPR dCAS9 activation (CRISPRa) targeting of an RA associated variant. The region around an associated RA variant (rs9603616) on chromosome 13 intronic of the COG6 genewas targeted in the HEK2937T cell line, using p300 as the activator. **a**, Fold change (qPCR) affect on FOXO1 gene expression compared to negative control. **b**, Fold change (qPCR) affect on COG6 gene expression compared to negative control.

## Discussion

We have generated a unique, high quality, high value resource, correlating a range of dynamic data, to inform the assignment of regulatory regions to genes. This analysis adds to the growing evidence that the relationship between enhancers and promoters is complex, that interactions are more strongly correlated within a distance of 200 kb, and that they are mostly constrained within TADs.

Using CD4+ T-cells, stimulated with anti-CD3/anti-CD28, we analyse ATAC-seq, RNA-seq, Hi-C and CHi-C over 24 hours. Time points were selected to understand how chromatin changes in the very early stages after stimulation, and how GWAS associated variants for autoimmune diseases, in particular RA, map to these chromatin dynamics. We show that the conformation of the DNA at a higher structural level of A/B compartments^28^ and TADs^29^ remains relatively constant throughout the stimulation time course. In contrast, DNA interactions at the level of discrete contacts, for example between open chromatin and gene promoters, is highly dynamic post stimulation, with over 30% of individual interactions showing some degree of change over 24 hours; however only a minority of these changes are correlated with a change in expression. These results suggest ATAC-seq peaks are associated with 2 CHi-C interactions on average, with each gene making contact with 7.7 ATAC-seq peaks on average. Although this correlation can occur over large distances (up to 5 Mb in our data), it is strongly enriched within 200 kb. We also demonstrate how subtle changes in both ATAC-seq and interaction intensity can have more marked effects on gene expression. These data therefore suggest that, when assigning GWAS variants to putative causal genes, all genes within 200 kb, all within the TAD structure, and multiple causal genes, should be considered as candidates for functional validation. In addition, as a proof of principle, we have confirmed one long range (~1 Mb) gene target using CRISPR/Cas9. Marks for active enhancers (H3K27ac and H3K4me1) are highly enriched in both all ATAC-seq peaks and those interacting with a gene promoter. However, since CTCF sites are also enriched under these ATAC-seq peaks, this does not exclude the possibility that these are involved in the dynamics of the data.

We also implicate genes in RA associated loci not previously highlighted as likely causal from GWAS, most notably MYC and FOXO1. MYC is a proto-oncogene transcription factor, involved in pro-proliferative pathways, highly expressed in a wide range of cancers. It has long been known that this gene is expressed in the RA synovium^30,31^, potentially playing a role in the invasiveness of these cells. More notably for this study, it has recently been demonstrated how a CD4+ T-cell subset from RA patients demonstrates higher autophagy, and that MYC is a central regulator of this pathway. Here it was suggested that autophagy could contribute to the survival of inflammatory T cells in patients, particularly a pathogenic-like lymphocyte (CPL) subset, found in inflamed joints and associated with disease activity. Similarly FOXO1 has long been established to be downregulated in both synovium and blood from RA patients, and is also correlated with disease activity^26^. FOXO1 is a transcription factor, thought to play a role in apoptosis and cell cycle regulation, where reduced expression in RA is suggested to have a role in the accumulation of fibroblasts in the disease synovium^26^. In this study, for both these genes with established biological mechanisms and expression patterns relevant to RA, we have demonstrated how genetic variants that lead to an increased risk of developing disease are physically linked and correlated with gene expression, providing evidence that these genes may be causal instigators in disease, and not simply on the pathways that are dysregulated in disease.

Our results indicate that: (i) both DNA activity and interaction intensity are independently important in the regulation of genes; and (ii) since a minority of interactions correlate with gene expression, simply assigning target genes by interactions is too simplistic. Instead, other methods, such as the simultaneous measurement of DNA activity and expression data followed by CRISPR experimental validation, are required to confidently assign genes to GWAS implicated loci. Finally, we confirm that subtle changes in interaction intensity are correlated with much larger changes to gene expression. In combination these findings have important implications for fully exploiting GWAS data, assigning causal SNPs, genes, cell types and mechanism to trait associated loci, on the pathway to translating these findings into clinical benefit.

## Methods

### Isolation of CD4+ T-cells and stimulation time course

Primary human CD4+ T-cells were collected from three healthy individuals with informed consent and with ethical approval (Nat Rep 99/8/84). Samples were isolated from PBMCs using an EasySep T-cell isolation kit (StemCell), plated in 6-well plates then stimulated with anti-CD3/anti-CD28 Dynabeads (Life Technologies) over a period of 24-hours, with samples removed at the appropriate time point and processed according to the experiment the cells would be used for. Unstimulated samples were also prepared (t=0 sample). For Hi-C experiments, CD4+ T-cells were harvested and fixed in formaldehyde, samples for RNA-seq (5 × 10^6^ CD4+ T-cells) were stored in RNA CellProtect reagent before extraction, and samples for ATAC-seq (50,000 cells) were processed immediately. ATAC-seq samples from three individuals were taken at time 0 mins, 20 mins, 1 hr, 2 hrs, 4 hrs and 24 hrs. Two pooled nuclear RNA-seq replicates were taken at the same time points as ATAC-seq samples. Two pooled Hi-C and CHi-C replicates were taken at 0 min, 20 min, 1 hr and 4 hrs and one sample for Hi-C and CHi-C was taken at 24 hrs, respectively.

### Library generation for CHi-C and Hi-C

To generate libraries for Hi-C experiments, 8-10 × 10^6^ CD4+ T-cells were harvested at the appropriate time point and formaldehyde crosslinking carried out as described in Belton et al^32^. Cells were washed in DMEM without serum then crosslinked with 2% formaldehyde for 10 minutes at room temperature. The crosslinking reaction was quenched by adding cold 1M glycine to a final concentration of 0.125M for five minutes at room temperature, followed by 15 minutes on ice. Crosslinked cells were washed in ice cold PBS, the supernatant discarded and the pellets flash-frozen on dry ice and stored at −80°C.

Hi-C libraries were prepared from fixed CD4+ T-cells from three individuals which were pooled at the lysis stage to give ~30 million cells. Cells were thawed on ice and re-suspended in 50ml freshly prepared ice-cold lysis buffer (10mM Tris-HCl pH 8, 10mM NaCl, 0.2% Igepal CA-630, one protease inhibitor cocktail tablet). Cells were lysed on ice for a total of 30 min, with 2×10 strokes of a Dounce homogeniser 5 min apart. Following lysis, the nuclei were pelleted and washed with 1.25xNEB Buffer 2 then re-suspended in 1.25xNEB Buffer 2 to make aliquots of 5-6×10^6^ cells for digestion. Following lysis, libraries were digested using HindIII then prepared as described in van Berkum et al^33^ with modifications described in Dryden et al^34^. Final library amplification was performed on multiple parallel reactions from libraries immobilised on Streptavidin beads using 8 cycles of PCR if the samples were to be used for CHi-C, or 6 cycles for Hi-C. Reactions were pooled post-PCR, purified using SPRI beads and the final libraries re-suspended in 30μl TLE. Library quality and quantity was assessed by Bioanalyzer and KAPA qPCR prior to sequencing on an Illumina HiSeq2500 generating 100bp paired-end reads (Babraham sequencing facility).

### Solution hybridisation capture of Hi-C library

Pre-CHi-C libraries corresponding to 750ng were concentrated in a Speedvac then re-suspended in 3.4#x03BC;l water. Hybridisation of SureSelect custom capture libraries to Hi-C libraries was carried out using Agilent SureSelectXT reagents and protocols. Post-capture amplification was carried out using 8 cycles of PCR from streptavidin beads in multiple parallel reactions, then pooled and purified using SPRI beads. Library quality and quantity was assessed by Bioanalyzer and KAPA qPCR prior to sequencing on an Illumina HiSeq2500 generating 100 bp paired-end reads (Babraham sequencing facility).

### Defining regions of association for bait design

All independent lead disease-associated SNPs for RA were selected from both the fine-mapped Immunochip study^20^ and a *trans*-ethnic GWAS meta-analysis^21^. This resulted in a total of 138 distinct variants associated with RA after exclusion of *HLA-*associated SNPs. Associated regions were defined by selecting all SNPs in LD with the lead disease-associated SNP (*r*^2^>=0.8; 1000 Genomes phase 3 EUR samples; May 2013). In addition to the SNP associations, credible SNP set regions were defined for the Immunochip array at a 95% confidence level.

### Target Enrichment Design

Capture oligos (120 bp; 25-65% GC, <3 unknown (N) bases) were designed to selected gene promoters (defined as the restriction fragments covering at least 500bp 5’ of the transcription start site (TSS)) using a custom Perl script within 400 bp but as close as possible to each end of the targeted HindIII restriction fragments and submitted to the Agilent eArray software (Agilent) for manufacture. Genes were selected as follows: all genes within 1Mb upstream and downstream of associated RA SNPs from Eyre *et al*^20^ and Okada *et al*^21^ as previously described; all gene promoters showing evidence of interacting with an associated region in our previous CHi-C study using GM12878 and Jurkat cell lines; all genes contained within the KEGG pathways for “NF-kappa B signalling”, “Antigen processing and presentation”, “Toll-like receptor signalling”, “T cell receptor signalling” and “Rheumatoid arthritis”; all genes showing differential expression in CD4+ T-cells after stimulation with anti-CD3/anti-CD28; all genes from Ye *et al*^4^ within the ‘Early induced’, ‘Intermediate induced I’ and ‘Intermediate induced II’ categories; and all genes from the Ye *et al*^4^ NanoString panel. Additionally control regions targeting the HBA, HOXA and MYC loci were included for quality control purposes.

### Library generation for RNA-seq

Nuclear RNA-seq was used to quantify nascent transcription to determine changes through time. Five million CD4+ T-cells were harvested, stored in Qiagen RNAprotect solution and the nuclear RNA isolated using a Qiagen RNeasy kit and quantified. Samples were either pooled in equal amounts (same individuals as for Hi-C to create matched samples), or processed individually to give duplicate samples. Libraries for RNA-seq were prepared using the NEB Next Ultra Directional RNA-seq reagents and protocol using 100ng of nuclear RNA as Input. Each library was sequenced on half a lane of an Illumina HiSeq2500 generating 100bp paired-end reads (Babraham sequencing facility).

### Library generation for ATAC-seq

ATAC-seq libraries were generated from 50,000 CD4+ T-cells from three individual samples using the protocol detailed in Buenrostro et al^35^ using the Illumina Nextera DNA Sample Preparation Kit. Each library was sequenced on half a lane of an Illumina HiSeq2500 generating 100 bp paired-end reads (Babraham sequencing facility).

### Hi-C data processing

Hi-C data were mapped to GRCh38 by HiCUP^36^. The maximum and minimum di-tag lengths were set to 800 and 150, respectively. HOMER^37^ Hi-C protocol was applied to Hi-C bam file and normalized Hi-C matrices were generated by analyzeHiC command from HOMER with resolution of 40,000bp (analyzeHiC –res 40000 –balance). TADs were generated by the command findTADsAndLoops.pl (-res 40000). A/B compartments were generated by runPCA.pl (-res 40000) followed by findHiCCompartments.pl with the default parameters to generate compartments A and –opp parameters to generate compartments B.

### CHi-C data processing

CHi-C data were mapped to GRCh38 by HiCUP. The maximum and minimum di-tag lengths were set to 800 and 150, respectively. CHiCAGO^38^ was applied to each bam file with the CHiCAGO score set to 0. Counts data for each interaction were extracted from the .rds files generated by CHiCAGO. Time course interactions were concatenated. Those interactions with at least one time point having CHiCAGO score over 5 were kept. Bait-to-bait interactions were registered as two interactions with either side defined as ‘bait’ or ‘otherEnd’.

### ATAC-seq data processing

Individual ATAC-seq reads data were mapped to GRCh38 by Bowtie2^39^ (with option -x 2000) and reads with length less than 30 were filtered by SAMtools^40^. Duplications were removed by Picard (https://broadinstitute.github.io/picard/). The three replicated bam files at each time point were merged by SAMtools. MACS2^41^ was applied on each merged bam file to call peaks (with option -- nomodel --extsize 200 --shift 100). Peaks generated from each time point were merged by Diffbind^42^ with default parameter to form the time course profile for ATAC-seq peaks.

### RNA-seq data processing

RNA-seq data were mapped to GRCh38 by STAR^13^ with default parameters. Counts data for exons and introns were generated by DEXSeq^43^. Individual counts data from each time point were combined to form the time course gene expression data. Exons and introns counts data were summed to get the gene expression data for each gene at each time point, respectively. Genes with the sum of counts data across the six time points less than 10 were removed in each replicate. Only genes that have expressions in both replicates were kept. 18,162 genes remained after this processing.

### Linking CHi-C, ATAC-seq and RNA-seq time course data

CHi-C time course data were linked to RNA-seq time course data with baits design specifying the mapping between baits and genes. ATAC-seq peaks residing at an otherEnd fragment were correlated with CHi-C interactions originating from that specific otherEnd fragment to different baits. Averaged data from replicates were used in correlation analysis. Pearson correlation coefficients between the connected CHi-C, gene and ATAC-seq time course data were calculated, respectively. Background random correlation tests were carried out by randomly picking up relevant time course data within the targeted dataset without any restrictions and calculating their Pearson correlation coefficients accordingly.

### Gaussian process test for dynamic time course data

Time course data were fitted by a Gaussian process regression model^16^ with a Radial Basis Function (RBF) kernel plus a white noise kernel (dynamic model) and a pure white noise kernel (static model), respectively. BIC was calculated for the dynamic model and flat model, respectively.

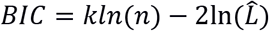

where k is the number of parameters used in the specified model, n is the sample size and 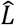 is the maximized likelihood for the model. Models with smaller BIC are favoured for each time course profile. Those with smaller BICs in dynamic models were classified as time-varying. A more stringent χ^2^ test with degree of freedom (df) of 1 was also applied to the Loglikelihood Ratio (LR) statistics, with 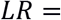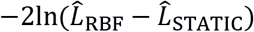, where 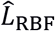 and 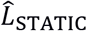 are the maximized likelihoods for the Gaussian process model and a static model, respectively. A p value of 0.05 was deemed significant.

### ATAC-seq data clustering and MOTIF searching

A more inclusive threshold of LR<-1 was applied to ATAC-seq peaks prior to clustering, which leaves 16% (12,215/74,583) ATAC-seq dynamical peaks, among which 9,680 were outside promoter regions ([+500bp,−1000bp] around genes). These ATAC-seq peaks were clustered using a Gaussian Process mixture model^44^. MOTIFs for each cluster were searched by findMotifGenome.pl (-mask –len 5,6,7,8,9,10,11,12 –size given) from HOMER with remaining peak data being used as background data.

### Construction of 99% credible SNP sets for RA loci

We considered 80 loci attaining genome-wide significance for RA in the European ancestry component of the most recently published trans-ethnic GWAS meta-analysis^21^, after excluding the MHC. For each locus, we calculated the reciprocal of an approximate Bayes’ factor in favour of association for each SNP by Wakefield’s approach^45^, given by

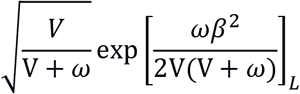

where β and V denote the estimated log odds ratio (log-OR) and corresponding variance from the European ancestry component of the meta-analysis. The parameter ω denotes the prior variance in allelic effects, taken here to be 0.04. The posterior probability of causality for the SNP is then obtained by dividing the Bayes’ factor by the total of Bayes’ factors for all SNPs across the locus. The 99% credible set for each locus was then constructed by: (i) ranking all SNPs according to their Bayes’ factor; and (ii) including ranked SNPs until their cumulative posterior probability of causality attained or exceeded 0.99.

### Overlap between ATAC-seq peaks and autoimmune GWAS loci

Autoimmune GWAS loci (defined by NCIT:C2889, OBI:OBI 1110054, OMIM:613551, OMIM: 615952, OMIM: 617006, SNOMEDCT:85828009) were downloaded from the GWAS catalog (https://www.ebi.ac.uk/gwas/efotraits/EFO_0005140, accessed July 2018) and SNPs with r^2^ ≥0.8 identified using PLINK v1.07. These were then overlapped with ATAC-seq peaks linked to CHi-C genes, using bedtools v2.21.0, to identify regions of open chromatin containing an autoimmune GWAS SNP SNP or highly correlated variant.

### RA heritability and enrichment

We performed RA SNP enrichment and heritability estimates in the ATAC peaks identified throughout the time course. This was performed using partitioned heritability analysis from the LD score regression software ^46^ and European ancestry data from the latest RA GWAS meta-analysis^21^. Briefly, the heritability of RA and SNP enrichments are computed in partitioned sections of the genome, in this case, the ATAC-seq peaks at each timepoint.

### CRISPR activation using dCas9-p300

#### Cell lines

HEK293T cells (clontech) were cultured in high glucose-containing Dulbecco’s modified Eagle’s medium (DMEM; Sigma) supplemented with 10% FBS and 1% penicillin streptomycin at 37oC/5% C02 and kept below passage 15.

#### Generation of the dCas9-p300 cell line and delivery of guides

HEK293T cells were first transduced lentivirally with the pLV-dCas9-p300-p2A-PuroR expression vector (addgene #83889) and were selected with 2ug/ml of puromycin and grown for a week before being banked as a cell line. A second round of lentiviral transduction was done to introduce the guide RNA (gRNA) using the vector pLK05.sgRNA.EFS.GFP (addgene #57822) and cells were doubly selected using both a maintenance selection of puromycin and sorted for the top 60% cells expressing GFP.

#### Guide RNAs

All guide RNAs were cloned into the guide delivery vector pLK05.sgRNA.EFS.GFP (addgene #57822) and are listed Table 1. A negative control guide Scr2 (AACAGTCGCGTTTGCGACT) is a scrambled guide sequence for comparison of gene expression that is not expected to target any known genes in the genome. A positive control guide IL1RN (CATCAAGTCAGCCATCAGC) was included, this is a guide directed to the transcription start site of the promoter of the *IL1RN* gene that has been previously shown to increase expression of the IL1RN gene substantially^47^. For the COG6/FOX01 locus three guides (TGGGGACTATCTAGCTGCT; AGGGCCTTATAATGTAGT; AGTCATCCTGGAGCACAGAGG) were pooled simultaneously in equimolar amounts to target the active enhancer marked by H3K27ac in proximity to the lead GWAS variant rs7993214.

### Lentivirus production

The day before transduction HEK293T cells were seeded at a density of 1E07 per transfer vector in 15cm plates in a volume of 20ml of DMEM 10% FBS without P/S. Each of the transfer vectors, together with packaging plasmids pmDLg/pRRE (#12251) and pRSV-REV (#12253) along with envelope plasmid pMD2.G (#12259), were combined to a total of 12ug at a ratio of 4:2:1:1, respectively in 2ml of serum free DMEM w/o phenol red.

PEI 1mg/ml was batch tested and added at a ratio of 6:1 PEI: DNA. The solution was briefly vortexed and incubated at room temperature for 15 minutes. Following this the solution was added dropwise to the cells. Flasks were rocked gently in a circular motion to distribute the precipitates, and then returned to the incubator. 24 hours later fresh growth medium was added of DMEM with 10% FBS and 1% P/S. The viral supernatant was collected 72 hours after transduction, cleared by centrifugation at 1500rpm for 5 minutes at 4⁰C and then passed through a 0.45um pore PVDF Millex-HV (Millipore). Lentivirus was aliquoted and stored at −80⁰C for future use.

### Transduction of HEK293T p300 cell line with the gRNAs

300,000 dCas9-p300-HEK 293T cells were plated onto 6 well plates in triplicate for each gRNA. 24 hours later the medium was changed to DMEM 10% FBS without penicillin streptomycin. 1ml of each gRNA generated lentivirus was added to each well of 300,000 HEK293T cells cultured in DMEM supplemented with 10% FBS in triplicate. 24 hours later the medium was changed to DMEM containing 10% FBS and 1% penicillin streptomycin. Cells were grown up for 5 days and then sorted for the top 60% of cells expressing GFP using flow cytometry.

### RNA extraction and qPCR

When confluent 2E06 cells were spun down at 400xg for 5 minutes and washed in PBS. RNA was extracted using the RNeasy mini kit (Qiagen) according to manufacturer’s instructions and the genomic DNA removal step was included. 100ng of RNA for each sample was used in a single RNA-to-Ct reaction (Thermofisher) to assay gene expression. Taqman assays FOX01 (Hs00231106_m1), COG6 (Hs01037401_m1) and IL1RN (hs00893626_m1) were used alongside housekeeping genes YWHAZ (Hs01122445_g1) and TBP (hs00427620_m1) for normalisation.

### Data analysis

Delta-delta CT analysis was carried out using the Scr2 generated dCas9-p300 HEK293T cells as the control and normalised against the YWHAZ and TBP housekeeping genes. The data was analysed in graph pad using one-way ANOVA.

## Supporting information

Supplementary Table 4 ALL RA loci

Supplementary Table 3 ALL Autoimmune loci

## Data availability

Raw sequencing data and processed counts data for ATAC-seq, RNA-seq, CHi-C and Hi-C that support the findings of this study have been deposited in National Center for Biotechnology Information’s Gene Expression Omnibus and are accessible through GEO Series accession number GSE138767 (https://www.ncbi.nlm.nih.gov/geo/query/acc.cgi?acc=GSE138767).

The full data is accessible using the following link:-http://epigenomegateway.wustl.edu/legacy/?genome=hg38&datahub=http://bartzabel.ls.manchester.ac.uk/worthingtonlab/functional_genomics/GSE138767/GSE138767.json

## Code availability

Scripts to reproduce the analysis and figures in this study are available on github https://github.com/ManchesterBioinference/IntegratingATAC-RNA-HiC/.

## Acknowledgements

We acknowledge support from Versus Arthritis (grant ref 21754) and MRC (grant ref MR/N00017X/1). P.M. is funded on Versus Arthritis Foundation fellowship (grant ref 21745). K.D. is funded on a Granville Hugh King foundation fellowship (grant ref 21146). The authors would like to acknowledge the assistance given by Research IT and the use of the Computational Shared Facility at the University of Manchester, UK. We acknowledge Servier Medical ART (https://smart.servier.com) for creating Fig. 1 in the main text.

## Author Contributions

M.R., S.E. and P.F. conceived the project. A.M. performed ATAC-seq and RNA-seq assays, Hi-C and CHi-C experiments. P.M. designed the Hi-C and CHi-C experiments. J.Y. analysed all the data, P.Z. X.G. and P.M. additionally analysed the data. A.A and K.D. designed and performed the CRISPR experiment. A.P.M. additionally analysed the ATAC-seq data. J.Y, A.P.M., M.R. and S.E. wrote the manuscript.

## Supplementary informations

### 1. RNA-seq data

Two pooled sample RNA-seq time courses were collected at times 0 mins, 20 mins, 1 hr, 2 hrs, 4 hrs and 24 hrs. Only samples with a RIN value of >9 were used to generate RNA-seq libraries. Reads were mapped to GRCh38 by STAR^1^ with default parameters. Counts data for exons and introns were generated using DEXSeq^2^. Exon and intron counts for the same genes show good correlations (Supplementary Fig. 1a-b). Gene counts data were generated by adding up exon and intron counts data for the same genes, which also correlated well between replicates (Supplementary Fig. 1c). Individual counts data from each time point were combined to form the time course gene expression data. Genes with the sum of counts data across the six time points less than 10 were removed in each replicate. Counts data from each replicate were merged to form the time course gene expression data used for clustering and correlation with other datasets. Only genes that showed expression in both replicates were kept. 22,126 genes were remained after these processing steps. Gene expression data were normalized by DESeq2^3^. Trajectories of principal component analysis (PCA) result of the two replicates on principal component 1 (PC1) and principal component 2 (PC2) is shown in Supplementary Fig. 1d, where the replicated data are grouped together for each time point, again illustrating good data quality. Gene expression data were compared to the data from Ye et al^4^, where similar experiments were carried out up to time 72 hrs. Time course profiles of the gene expression of the five categorized gene sets from Ye et al^4^, including ‘Early induced’, ‘Intermediate induced I’, ‘Intermediate induced II’, ‘Persistent repressed’ and “Late induced”, are shown in Supplementary Fig. 1e-i. Our data show similar patterns to those in Ye et al^4^.

**Supplementary Fig. 1.**
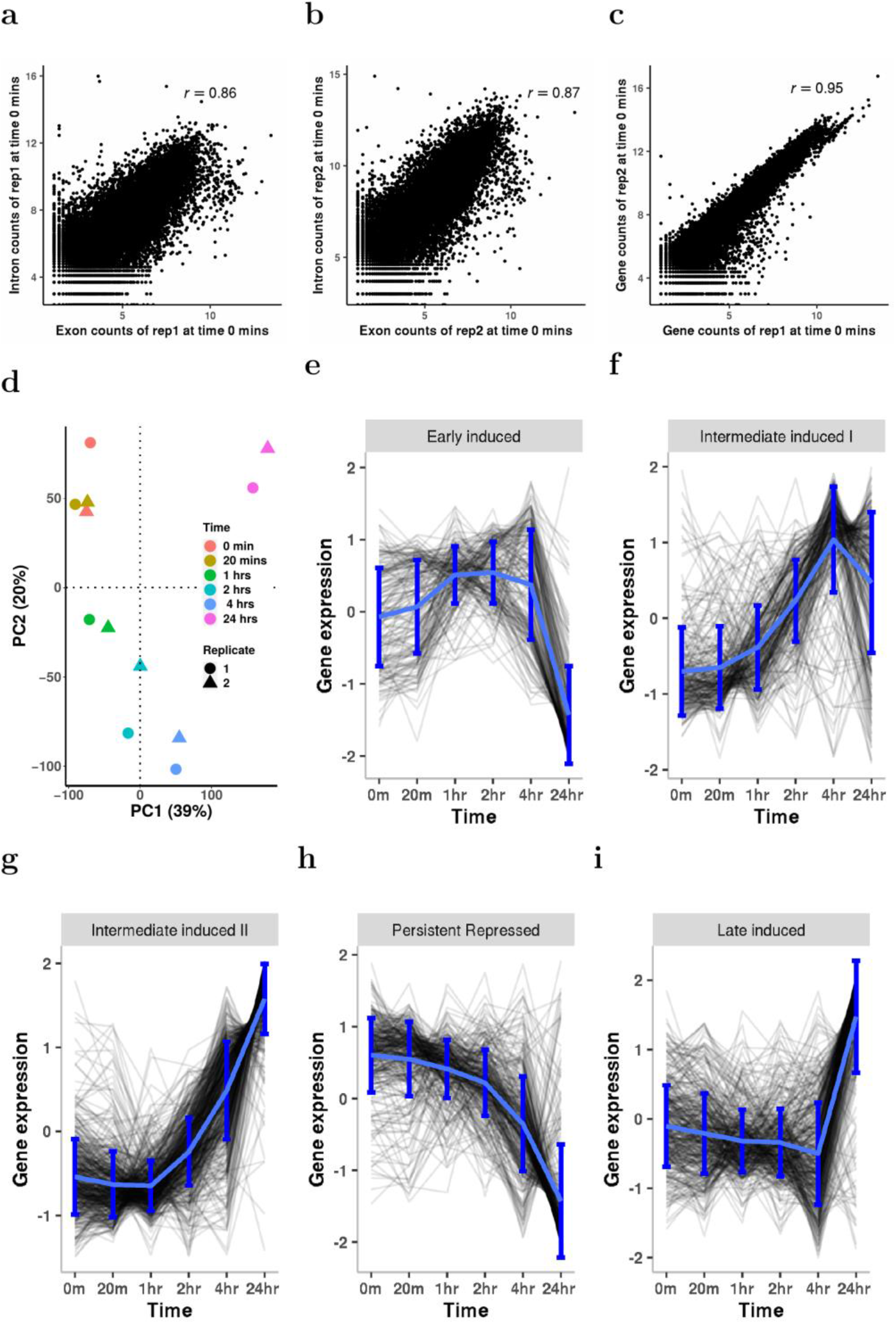
Illustration of RNA-seq time profiles. **a**, Scatter plot between natural logscaled exon and intron counts data of replicate 1 at time 0 mins. r is the Pearson correlation coefficient. **b**, Scatter plot between natural log scaled exon and intron counts data of replicate 2 at time 0 mins. **c**, Scatter plot between natural logscaled gene counts data of replicate 1 and replicate 2 at time 0 mins. **c**, PCA results of the two RNA-seq replicates used in this study. **e-i**, Time course profiles of gene sets as categorized in Ye et al^4^. Grey lines represent normalized gene expression and blue lines represent the mean of the data in each dataset. Errorbars are ± std of the data in each plot.

### 2. FACS analysis

We used an initial FACS panel (Supplementary Table 1) to determine our CD4+ T cell material was ~ 94% pure.

**Supplementary Table 1.** FACS panel for determining CD4+ T cell purity.

| Antibody | Wavelength | Dilution |
| --- | --- | --- |
| **TruStain FcX** |  | **(1:100)** |
| **CD3-APC Cy7** | **640 780/60** | **(1:50)** |
| **CD4-PerCPCy5.5** | **488 695/40** | **(1:100)** |
| **CD14-PE Cy7** | **488 780/60** | **(1:100)** |
| **CD19-BV605** | **405 610/20** | **(1:100)** |
| **CD8-BV785** | **405 780/60** | **(1:100)** |

### 3. Cell composition and effect of stimulation on CD4+ T cells

We performed absolute deconvolution using ABsolute Immune Signal (ABIS) deconvolution^5^ to investigate the degree of T memory and T effector cell composition (Supplementary Fig. 2a). To assess the amount of stimulation we explored markers of naivety, activation and cytokine expression and confirmed a similar degree of stimulation (Supplementary Fig. 2b).

**Supplementary Fig. 2.**
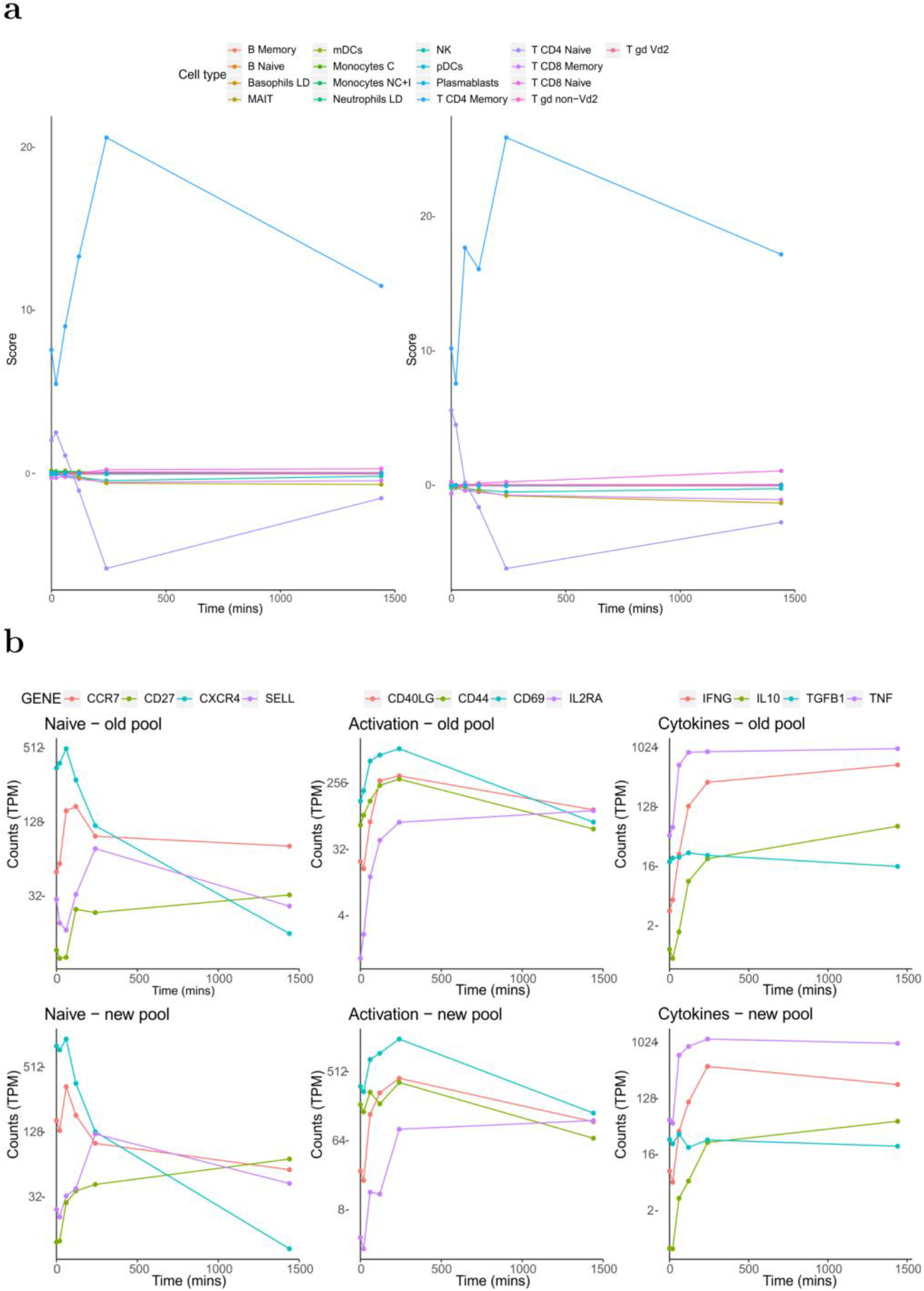
RNA-seq data used to determine initial CD4+ T cell composition (a) and the stimulation of cells (b) in the replicate pools.

### 4. ATAC-seq data

#### 4.1 ATAC-seq data

Three replicated ATAC-seq time series were collected at times 0 mins, 20 mins, 1 hr, 2 hrs, 4 hrs and 24 hrs. In total 18 ATAC-seq samples were sequenced. Sequenced raw reads were mapped to GRCh38 by Bowtie2^6^ (with option -x 2000) with reads mapping quality lower than 30 filtered out using SAMtools^7^ (with option -q30). Duplicates were removed by Picard (https://broadinstitute.github.io/picard/ with options MarkDuplicates REMOVE_DUPLICATES=TRUE). Less than 1.0% reads among these deduplicated reads were mitochondria reads (0.6% ± 0.3%) and were removed accordingly. The three replicated bam files at each time point were merged by SAMtools. Macs2^8^ was applied on each merged bam file to call peaks with default q value of 0.05 (with option --nomodel --extsize 200 --shift 100). Supplementary Fig. 3a shows the number of peaks identified for each time point and the number of peaks intersected between different time points. PCA was applied to these 18 ATAC-seqs and Supplementary Fig. 3e shows the distribution of these samples across PC1 and PC2. The results clearly illustrate substantial variability in the data. However, the time difference between samples is well captured by PC2. To align ATAC-seq datasets across time, the peaks generated from each time point were merged by Diffbind^9^ (with option minOverlap=1) and 76,359 peaks were obtained in the end with an average size of 488.32 bp. 74,583 peaks were retained with peak sizes within a 5-95 percentile range of 123 bp to 1414 bp (Supplementary Fig. 3c) after merging peaks across the six time points, with each peak appearing in at least one time point. ATAC-seq peaks were enriched for both marks of enhancer activity (ChromHMM, Supplementary Fig. 4a) and CTCF sites across all time points (Supplementary Figure 4b). ATAC-seq peaks were annotated using ChIPpeakAnno^10^ (Supplementary Fig 3d). Peaks that lie in the Promoter region (22.9%) were removed in the downstream analysis in order to focus on enhancer-associated peaks. Supplementary Fig3a shows the number of peaks identified for each time point and the number of peaks intersected between different time points. The FRIP (the Fraction of Reads In called Peaks) scores we obtained (Supplementary Fig. 3b) are comparable to those results from the original ATAC-seq method^11^. Supplementary Fig. 3f compares the peaks from our data and those peaks from Gate et al^12^. Due to different experiment setup, there are some discrepancies between the peaks from these two sources. However, the majority of peaks from our data are within the peaks from Gate et al^12^.

**Supplementary Fig. 3.**
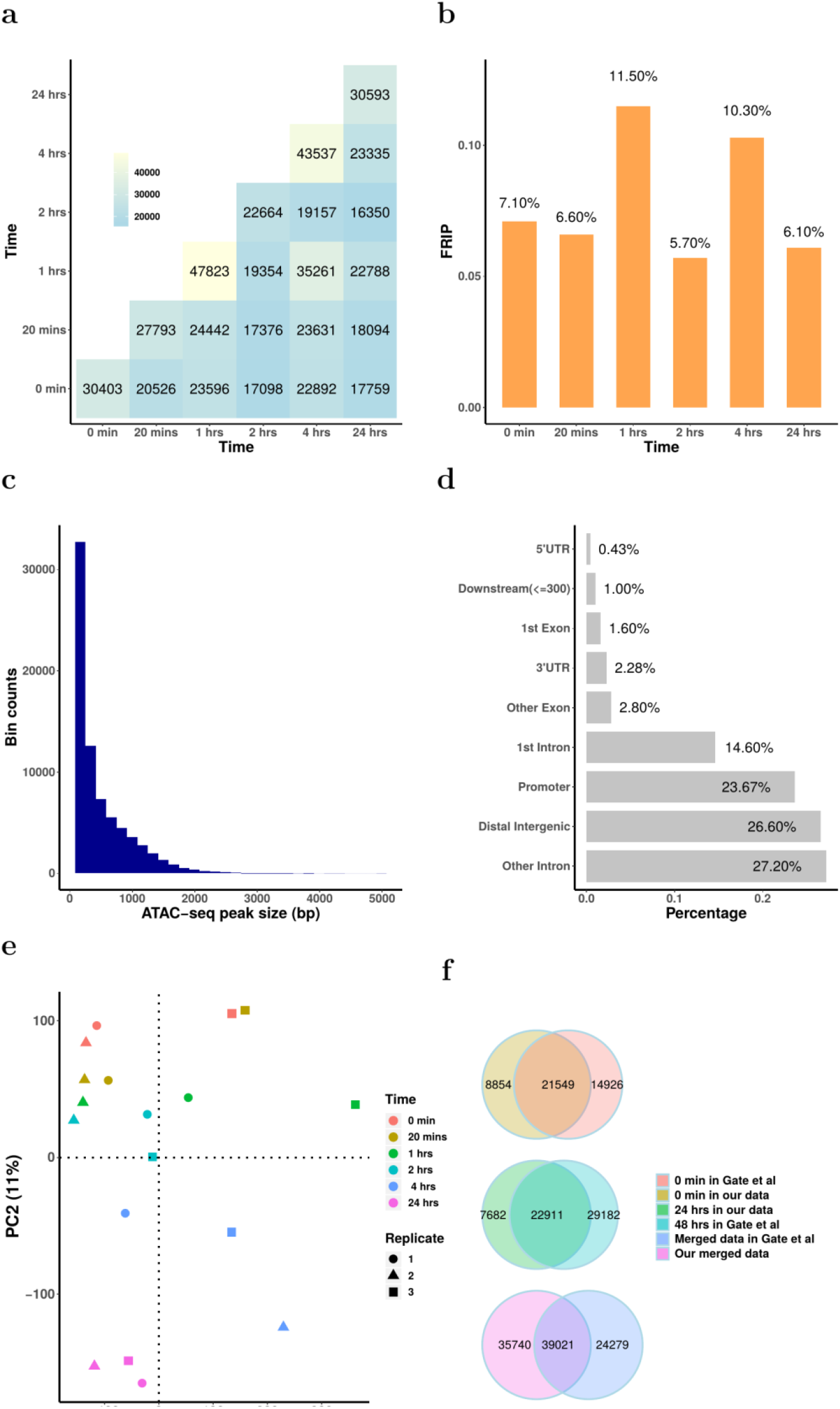
ATAC-seq peak quality check summary. **a**, Number of peaks at each time point and intersected number of peaks between different time points. **b**, FRIP scores for each time point. **c**, Histogram of ATAC-seq peak sizes. **d**, Bar plot of the annotations of the ATAC-seq peak data. **e**, Trajectories of PCA analysis of ATAC-seq replicates on PC1 and PC2. **f,** Venn diagram of the peaks from our study and peaks from Gate el al^12^.

#### 4.2 ATAC-seq peaks a proxy for enhancer activity or CTCF site, or both

We overlapped our ATAC-seq peaks with chromHMM chromatin state results from Roadmap epigenomics ^13,14^, we find that our ATAC peaks show an enrichment for enhancer and promoter states identified in CD4 T cells (Supplementary Figure 4a-d). We also note that there is an enrichment for CTCF binding sites in our ATAC peaks (Supplementary Figure 4e), particularly the ATAC-seq peaks around TAD boundaries (Supplementary Figure 4f).

In our analysis, enrichment was computed as follows, our ATAC-seq peaks were randomly shuffled 100 times, and overlapped with chromHMM chromatin state results. These shuffled overlaps were used as the baseline overlap. The empirical overlap was then used to calculated a fold change over the baseline overlap.

We note that while there remains an enrichment for both CTCF and enhancer/promoter states in both ATAC peak conditions across the timepoints, there is consistently more CTCF enrichment at TAD boundaries when compared to the ATAC peaks found inside TADs. From this we can conclude that ATAC peaks are better described as “open chromatin regions” as they are comprised of both enriched CTCF and enhancer/promoter activity.

**Supplementary Fig. 4.**
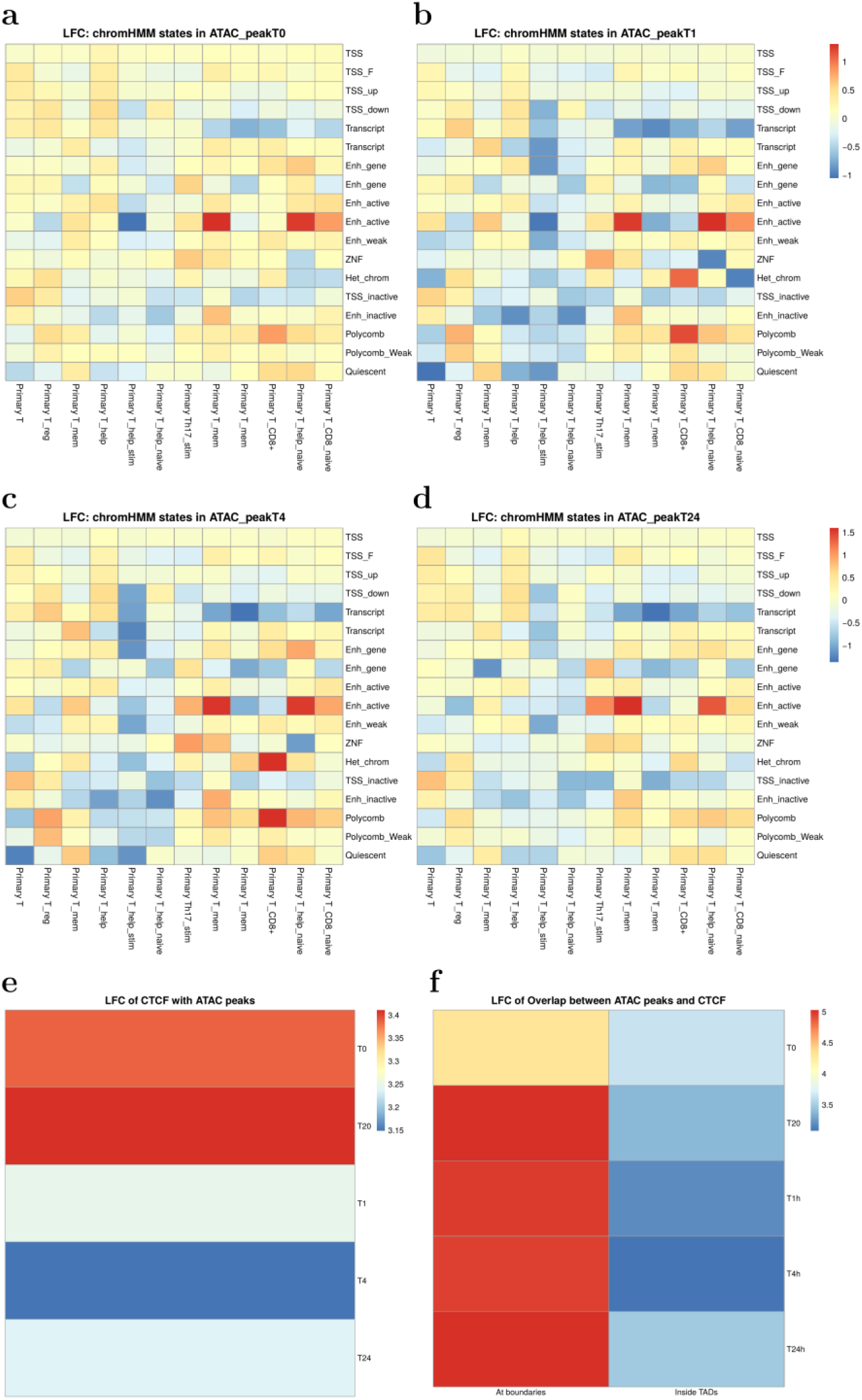
Demonstration of T cells chromHMM states enrichment at ATAC-seq peaks indentified at different timepoints. **a**, at time 0 min; **b**, at time 1 hrs; **c**, at time 4 hrs; **d**, at time 24 hrs. **e,** Enrichment of CTCF binding at all ATAC-peaks. **f,** Enrichment of CTCF peaks at TAD boundaries.

### 5. Capture Hi-C

A total of 7,081 baits were designed to capture 5,124 genes for CHi-C, among which 6,888 baits were successfully recovered with 97% on target. On average, 90.9±20.2 million unique di-tags were generated from two individuals for five experimental time points: 0 mins (unstimulated), and then 20 mins, 1 hr, 4 hrs and 24 hrs. One pooled sample Capture Hi-C (CHi-C) dataset was collected and sequenced at times: 0 mins, 20 mins, 1 hr and 4 hrs. Another pooled sample CHi-C dataset was sequenced at the same times as well as an additional 24 hr time-point. The CHi-C sequence data were mapped to GRCh38 using HiCUP^15^. The maximum and minimum di-tag lengths were set to 800 and 150, respectively. CHiCAGO^16^ was applied to each bam file with the CHiCAGO score set to 0. Counts data for each interaction were extracted from the.rds files generated by CHiCAGO. Interactions occurring at different time points (time 0 mins, 20 mins, 1 hr, 4 hrs and 24 hrs) were combined to create a complete set of all interactions and these were associated with counts for each time-point. Those interactions with at least one time point having CHiCAGO score over 5 were kept for further analysis. Bait-to-bait interactions are registered as two interactions with either side defined as “bait” or “otherEnd”. 271,398 interactions were generated this way, among which 17,196 interactions were trans-interactions and 254,202 interactions were cis-interactions. 6,888 baits and 121,656 otherEnds were involved in these interactions. CHi-C interactions occurring at either time 0 mins, time 4 hrs or both were extracted and compared to the interactions from Burren et al^17^. Venn diagrams of interactions originating from the two works under varied distance thresholds between bait and otherEnd fragments are shown in Supplementary Fig. 5a. It is clear that the closer the bait to otherEnd interactions are, the higher percentage of interactions are common between our experiments and experiments from Burren et al^17^. The top 24 clusters of the CHi-C interaction count time course data are illustrated in Supplementary Fig. 5b, showing varied and highly dynamic patterns of response.

**Supplementary Fig. 5.**
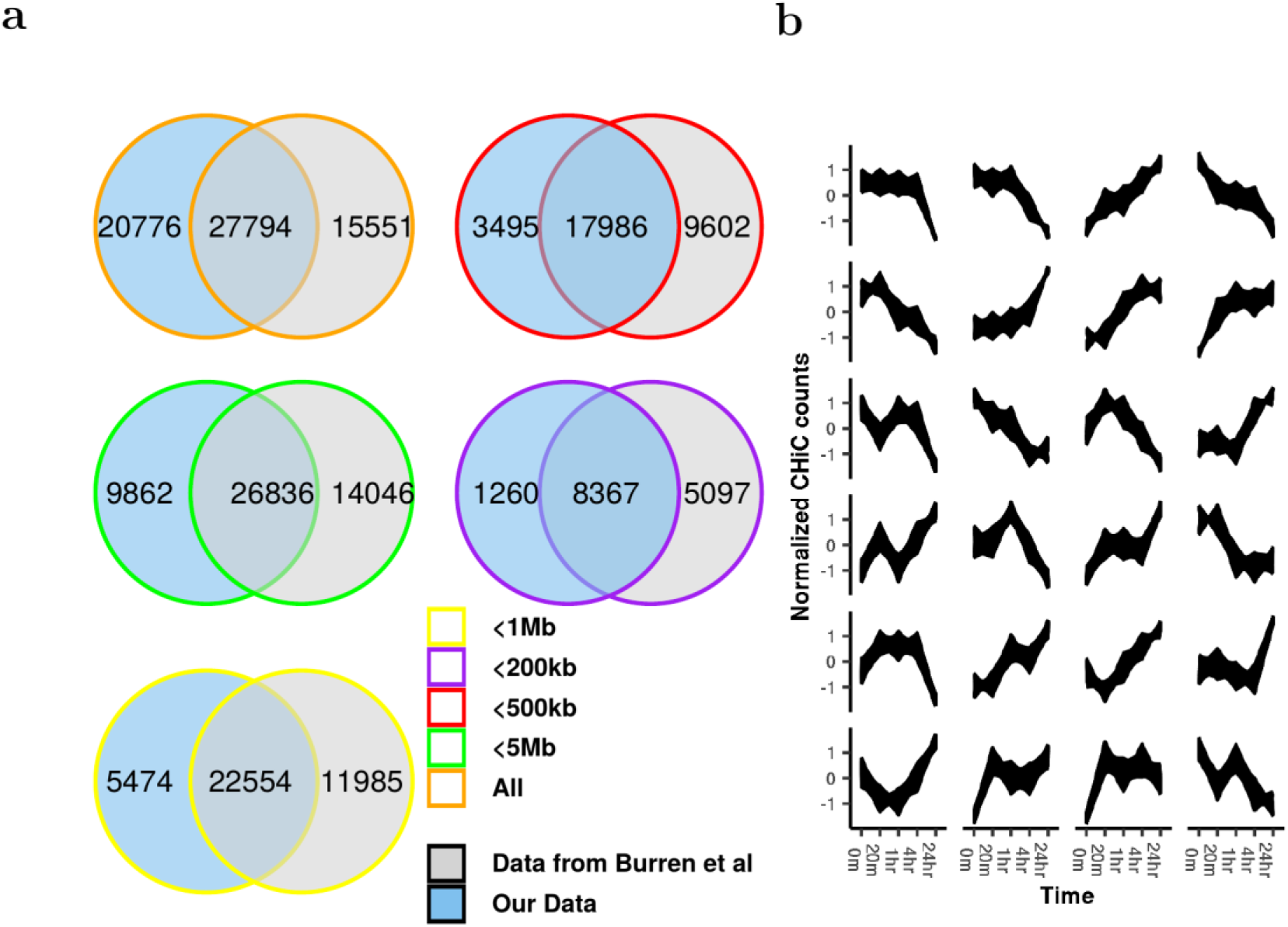
CHi-C quality check and time profile illustration. **a**, Venn diagram of the number of interactions interacted and remained between our data and data from Burren et al^17^ under different distance thresholds: “ All”, “<5Mb”, “<1Mb”, “<500kb”, “<200kb” representing range of distances to promoters considered. **b**, Largest 24 clusters of the time course profiles of CHi-C interactions within 5 Mb of promoters, using k-means clustering.

### 6. Hi-C data

#### 6.1 Hi-C data processing

One pooled sample Hi-C time course was collected at times 0 mins, 20 mins, 1 hr and 4 hrs. Another pooled sample Hi-C time course was collected at these times and an additional time of 24 hrs. Hi-C data were mapped to GRCh38 by HiCUP^15^ and then converted to HOMER^18^ format by scripts provided in HiCUP. The maximum and minimum di-tag lengths were set to 800 and 150, respectively. HOMER was applied the mapped Hi-C data (analyzeHiC –res 40000 -balance) and 1,230 distinct TADs were discovered across all 5 time points. Supplementary Fig. 6a shows fractions of the intersections of TADs across different time points within and between replicates. In the plot, intersections between row T_m_ and column T_n_ are defined by reciprocal 90% region overlap between the two TADs dataset at time T_m_ and T_n_. Fractions are then calculated by the number of intersected TADs between divided by the number of TADS at time point T_n_, respectively. A/B compartments were found by HOMER with command runPCA.pl (with option –res 40000) followed by findHICCompartments.pl to find A compartments or findHICCompartments.pl –opp to find B compartments, respectively. 1,136 A compartments and 1,266 B compartments were discovered. Supplementary Fig. 6b and 6c shows fractions of the A and B compartment across different time points within and between replicates. Here intersections are defined by reciprocal 90% region overlap between the two A/B compartment dataset at each time point. Fractions of intersections are calculated in the ways similar to that used for TADs.

#### 6.2 Changing A/B compartment overlap analysis

A/B compartments that changed over time were analysed for enrichment of (Lamina-associated domains) LADs ^19^ and gene classes (Supplementary Fig. 7). The log fold changes (LFC) were computed with the same method described above for ATAC-seq peaks. Chromosome positions of A/B compartments were converted to bed files and visualised in the UCSC browser^20^ (Supplementary Fig. 8).

**Supplementary Fig. 6.**
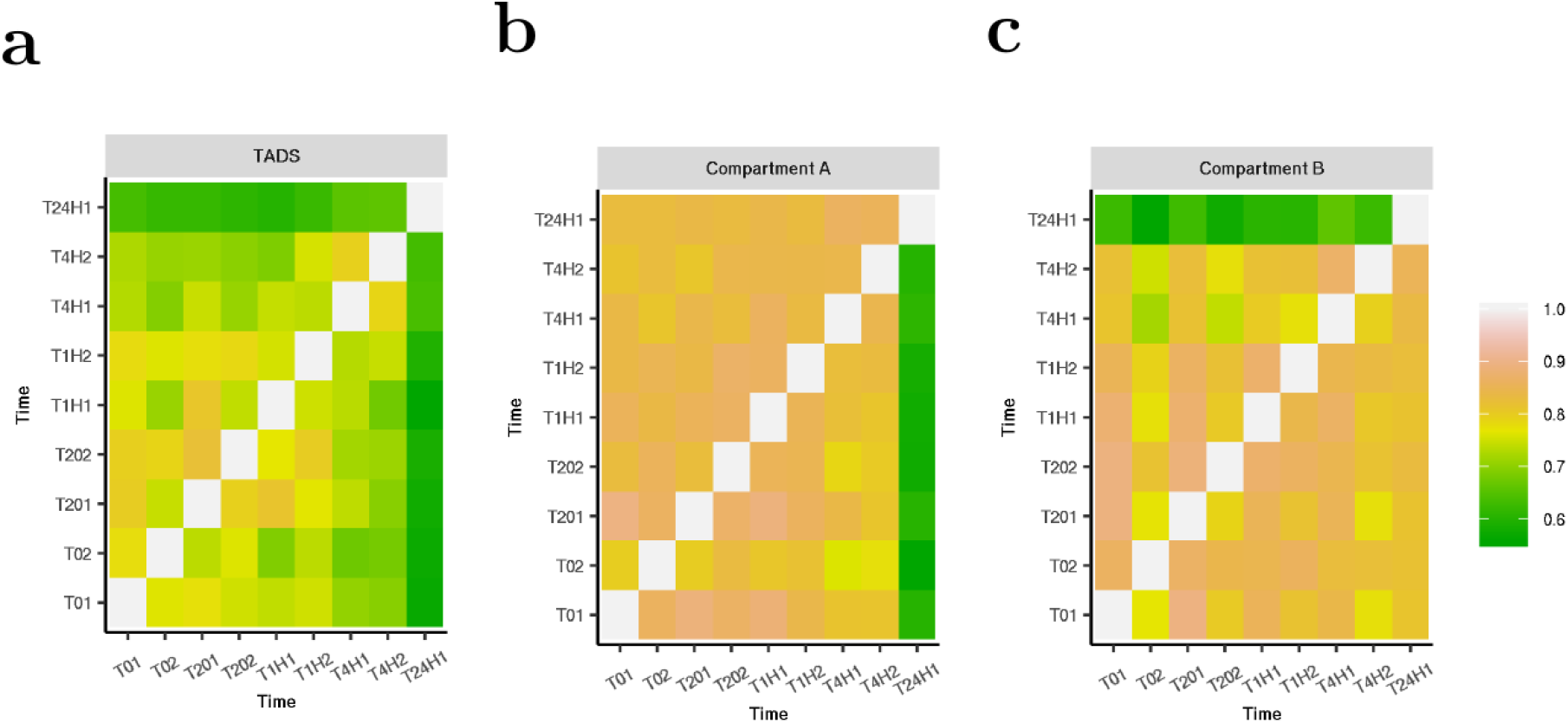
Fractions of TADS and AB compartments intersected. **a**, Fractions of intersected TADs with respect to the number of TADs at each time point in the horizontal axis. **b**, Fractions of intersected compartment As with respect to the number of compartment As at each time point in the horizontal axis. **c**, Fractions of intersected B compartments with respect to the number of compartment Bs at each time point in the horizontal axis. Here T01, T201, T1H1, T4H1, T24H1 are the time points for replicate 1 at time 0 min, 20 mins, 1 hrs, 4 hrs and 24 hrs, whereas T02, T202, T1H2, T4H2 are the time points for replicate 2 at time 0 min, 20 mins, 1 hrs and 4 hrs, respectively.

**Supplementary Fig. 7.**
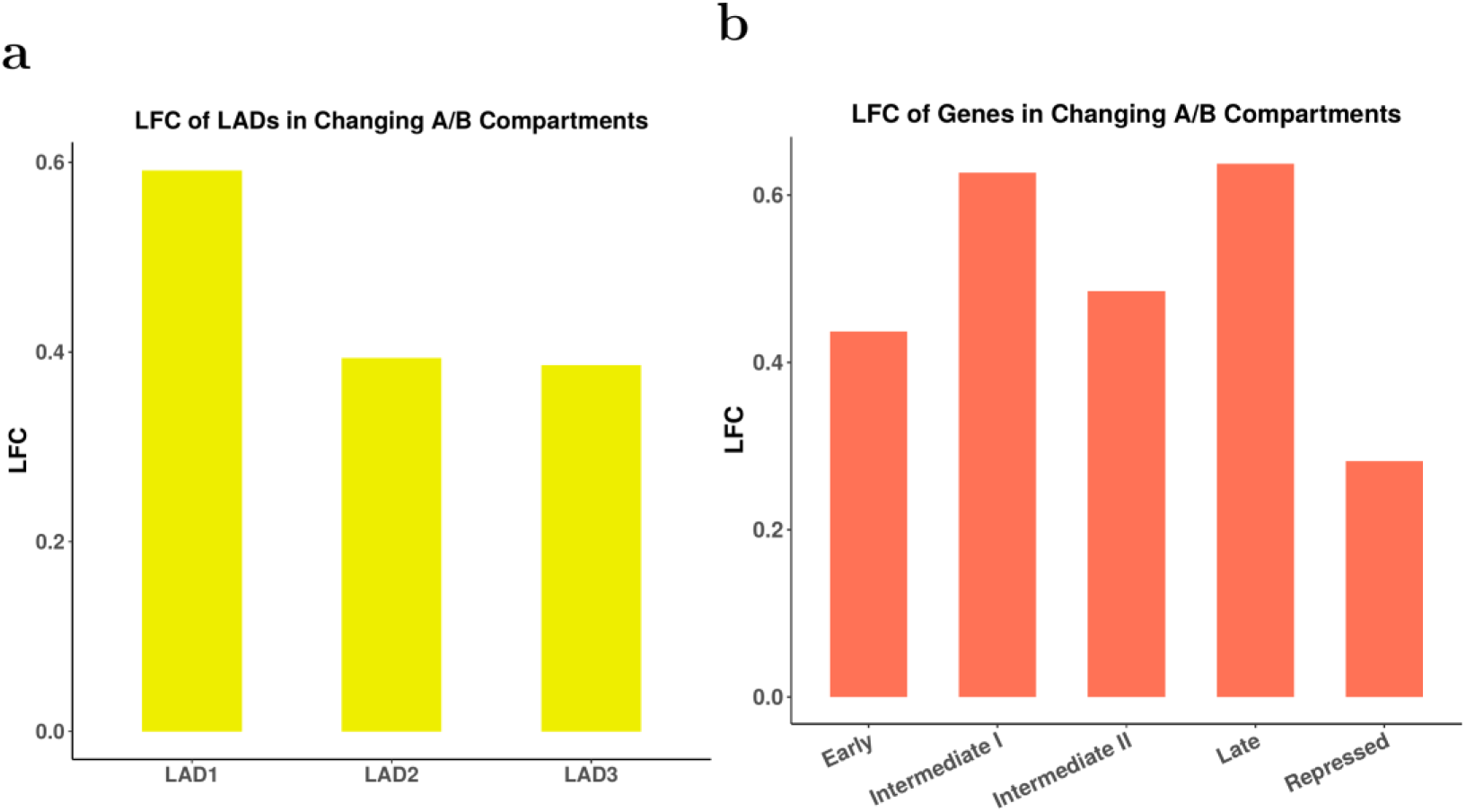
Enrichment analysis of changing A/B compartments over time. Log fold changes (LFC) of chromatin features overlaid with changing A/B boundaries, demonstrated an enrichment of both LADs (a) and expressed gene classes (b).

**Supplementary Fig. 8.**
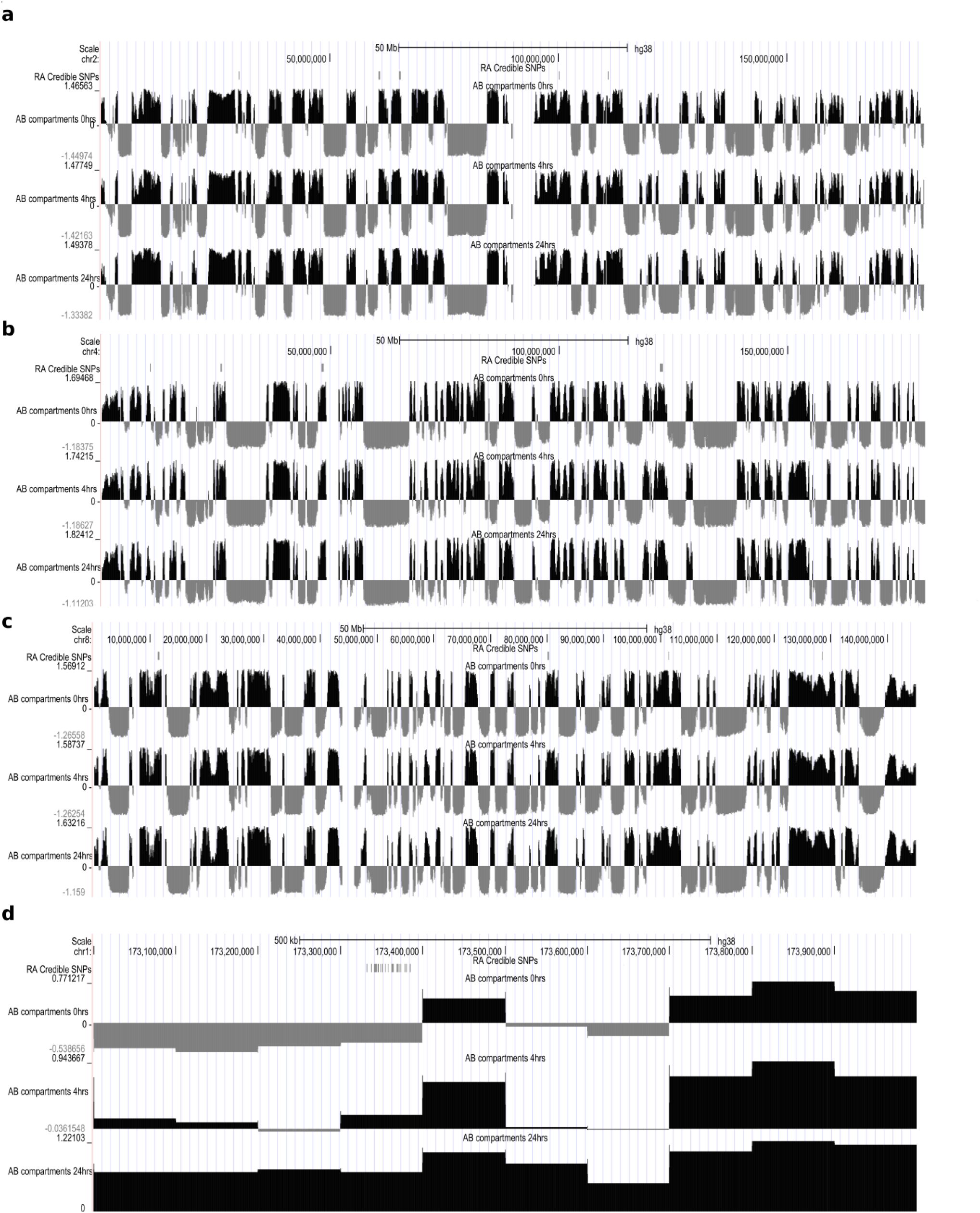
UCSC genome browser plots of chromatin structure, overlaid with epigenomic features and RA associated variants. **a-c**, Exemplar plots from chromosomes 2, 4 and 8 of A/B compartments (A compartments black; B compartments grey) called in at 3 time points (0, 4hrs and 24hrs) in duplicate T cells. **d**, A/B compartments on Chr1 around the TNFSF18 and TNFSF4 gene region. RA associated variants go from a largely inactive B region at 0hrs, through to an active A compartment at 24hrs.

### 7. ATAC-seq data clustering and MOTIF search

ATAC-seq data residing outside promoter regions, with loglikelihood ratio (LR, see main paper Methods) between a dynamic and static model over 1, were clustered using a hierarchical Gaussian Process mixture model^21^. MOTIFs for these clusters were searched by findMotifsGenome.pl from HOMER (-len 5,6,7,8,9,10,11,12,13 –size given) with ATAC-seq data with *BIC*_RBF_ > *BIC*_NOISE_ (static peaks) as background data. Clustered ATAC-seq peaks were intersected with ChIP-seq data for CTCF, NFAT1, JunB, NFKB^22^ and H3K27ac^14^ from CD4+ T cells. Z-scored data of these clustered ATAC-seq and associated CHi-C and gene expression data were examined and data over 0, equal 0 and less than 0 were labelled “+1”,”0” and “−1”, respectively, in order to assess the temporal change of the time over 24 hrs period. Supplementary Fig. 9a, 9b and 9c illustrate the temporal pattern change of clustered ATAC-seq, CHi-C and gene under ATAC-seq peaks with CTCF or H3K27ac binding only. In Supplementary Fig. 9d, CHi-C data associated with clustered ATAC-seq peaks were defined as “Gain”, “Lost” or “No change” based on if the count value difference between time 24 hrs and time 0 min is positive, negative or zero. Histograms of “Gain” or “Lost” on the six ATAC-seq clusters with ATAC-seq peaks binding with CTCF, JunB, NFAT1, NFKB and H3K27ac are illustrated.

**Supplementary Fig. 9.**
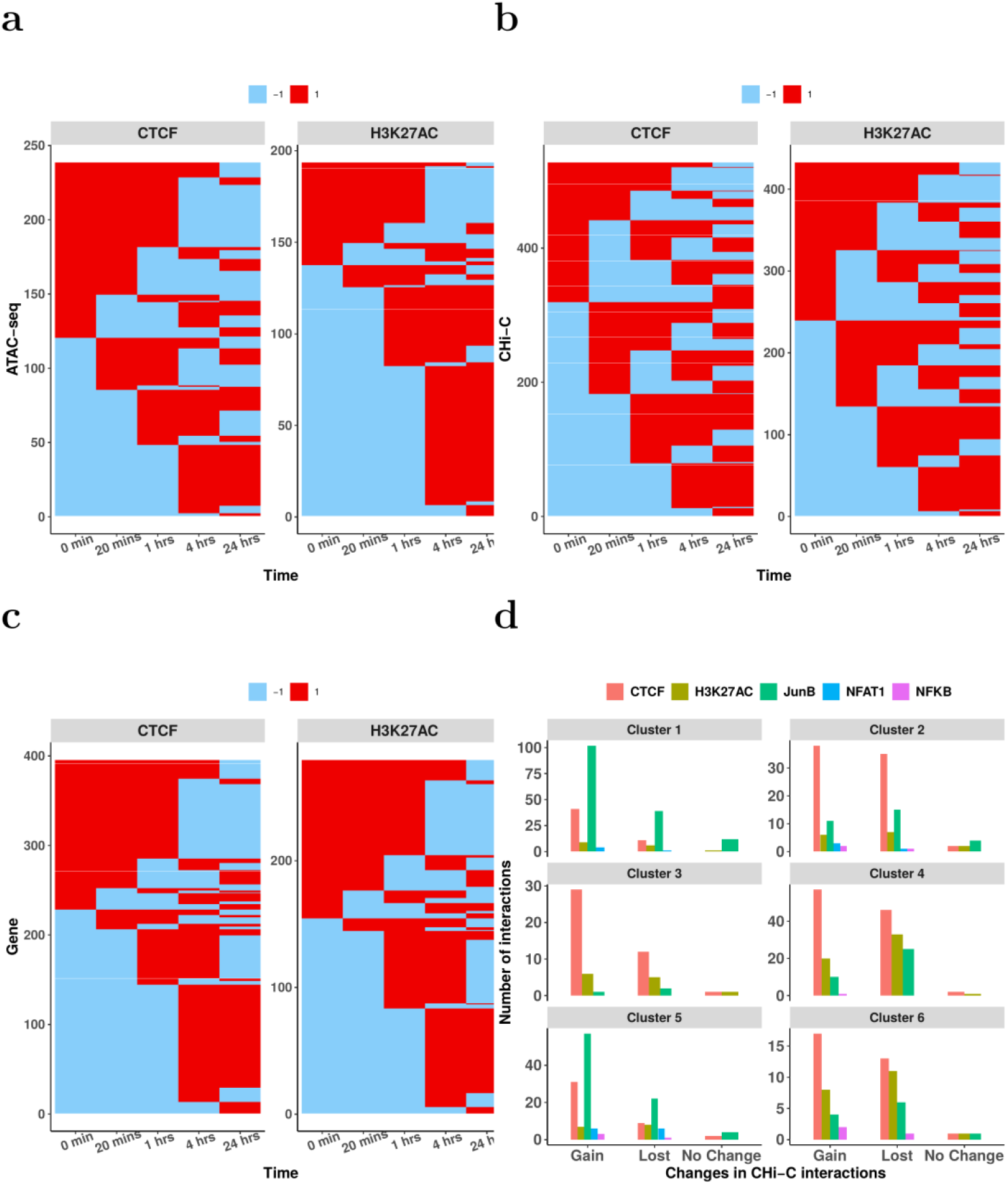
Temporal change of ATAC-seq, CHi-C and gene expression with different transcription factor binding. **a**, Comparisons of temporal change of ATAC-seq peaks with CTCF/H3K27ac binding. **b**, Comparison of temporal change of CHi-C interactions with CTCF/H3K27ac binding at associated ATAC-seq peaks. **c**, Comparison of temporal change of gene expressions with CTCF/H3K27ac binding at associated ATAC-seq peaks. **d**, Histograms of “Gain”/ “Lost”/ “No change” of CHi-C interactions under the six clustered ATAC-seq peaks binding with transcription factors including CTCF, H3K27ac, NFAT1, NFKB and JunB.

### 8. Correlations between ATAC-seq, CHi-C and gene

For ATAC-seq peaks sitting inside otherEnd fragments, the correlations between ATAC-seq time course, CHi-C time course and RNA-seq time course related to interacted baits are examined. Clusters between genes, CHi-C and ATAC-seq within the distance range of 200 kb of promoters under dynamical or stationary scenarios are shown in Supplementary Fig. 10a-f, respectively

**Supplementary Fig. 10.**
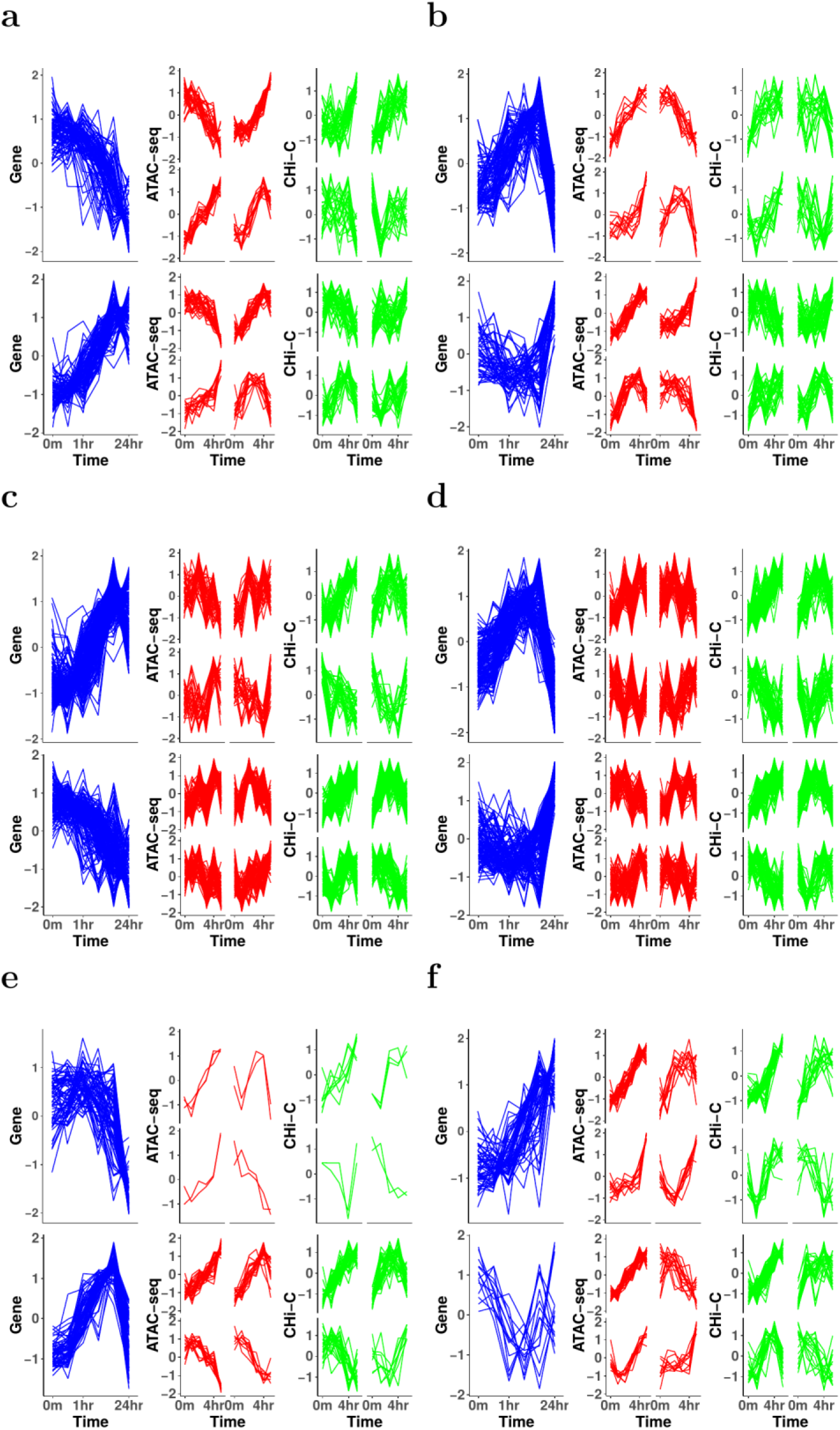
Clusters of gene expression, ATAC-seq counts and CHi-C data within 200 kb of promoters. **a,b**: Clusters with only ATAC-seq peaks are dynamical; **c,d**, Clusters with only CHi-C data are dynamical; **e,f**: Clusters with both ATAC-seq and CHi-C data are dynamical.

### 9. RA SNP and ATAC-seq peaks

#### 9.1 RA heritability and enrichment

We performed RA SNP enrichment and heritability estimates in the ATAC peaks identified throughout the time course, and have plotted the results below. This was performed using partitioned heritability analysis from the LD score regression software ^23^. Briefly, the heritability of RA and SNP enrichments are computed in partitioned sections of the genome, in this case, the ATAC-seq peaks at each timepoint. We observe an 5-30 fold enrichment of RA variants in the ATAC peaks, with a general trend of increasing enrichment post stimulation. RA SNP variants demonstrated an enrichment (5-30 fold) in open regions of chromatin, as expected, demonstrating the highest enrichment at 24 hours post stimulation (Supplementary Fig. 11).

**Supplementary Fig. 11.**
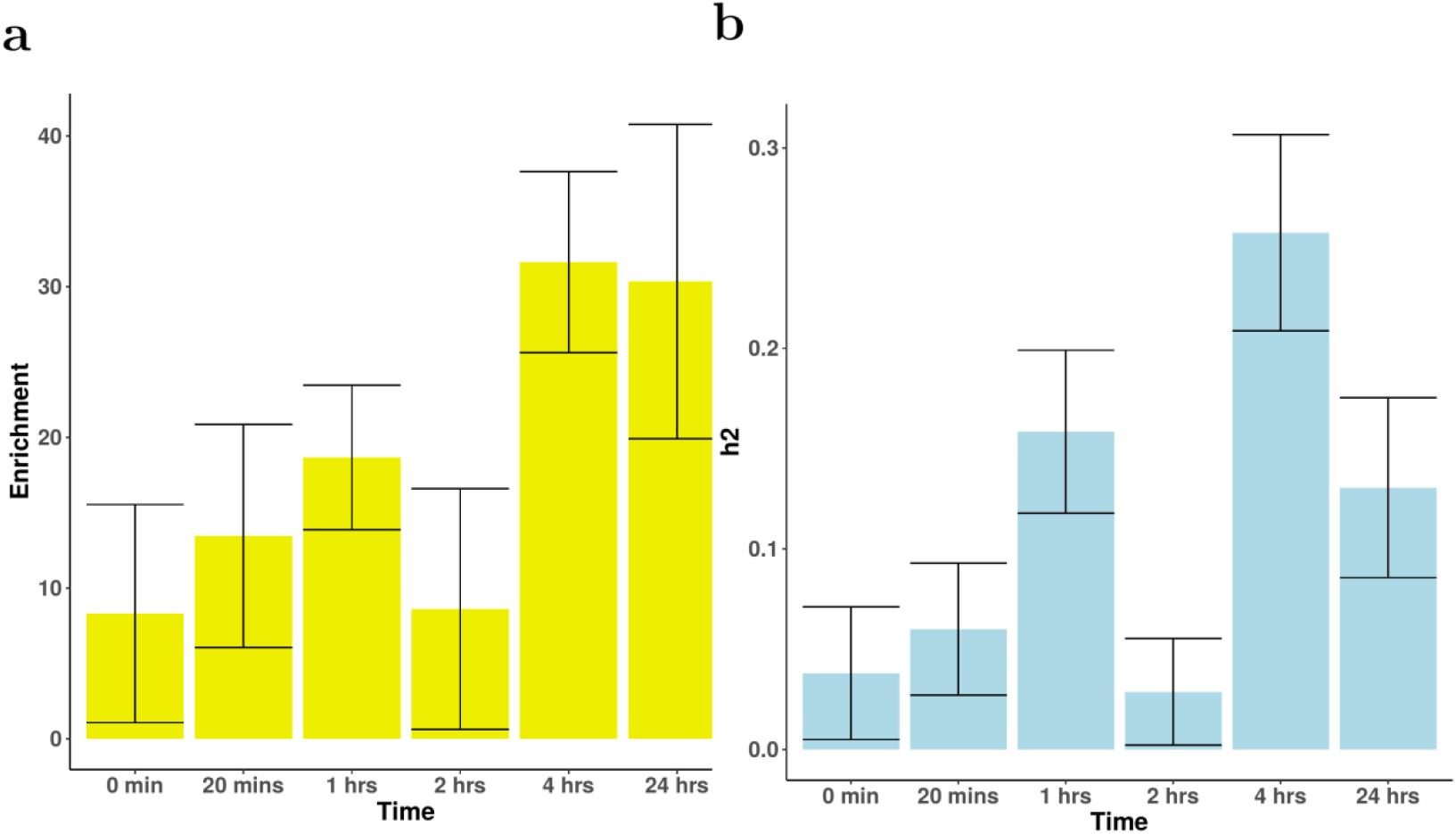
**a**, Enrichment of RA variants at ATAC-seq peaks at each time point. **b**, Partitioned heritability attributed to RA variants found within ATAC-seq peaks at different time points.

#### 9.2 Prioritisation of causal gene in RA loci

**Supplementary Fig. 12.**
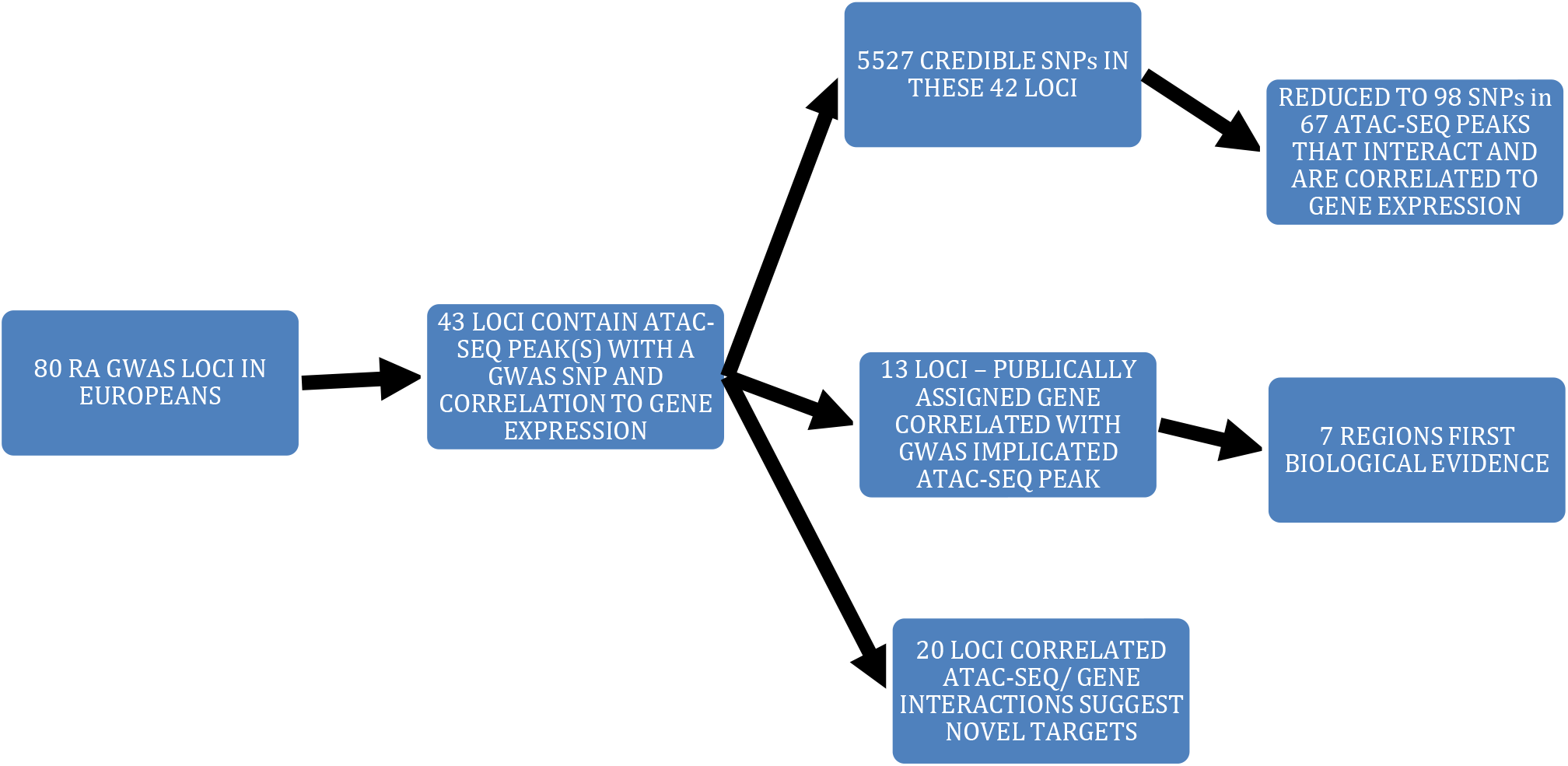
Schematic of pipeline for prioritisation of RA GWAS associated variants.

**Supplementary Fig. 13.**
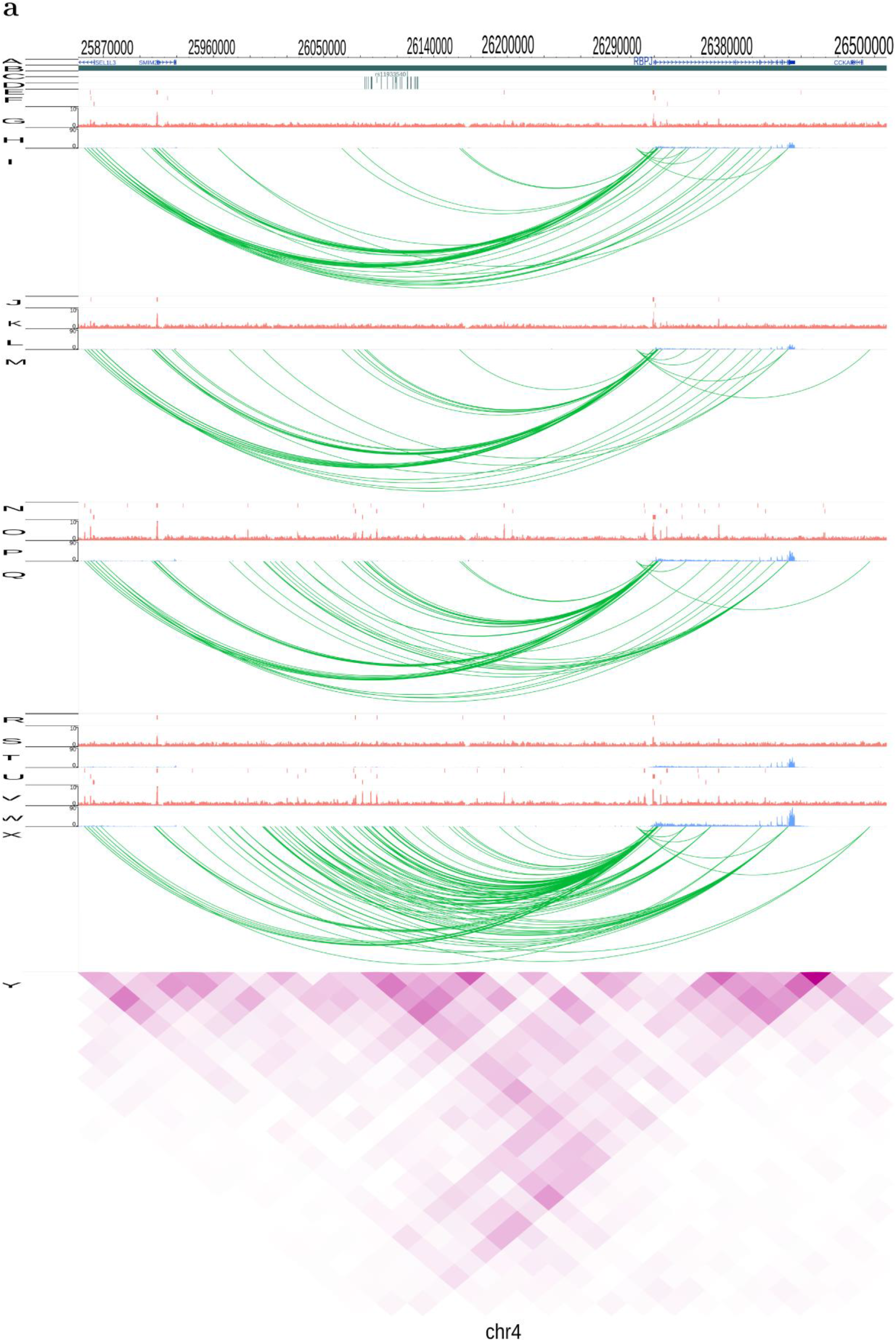
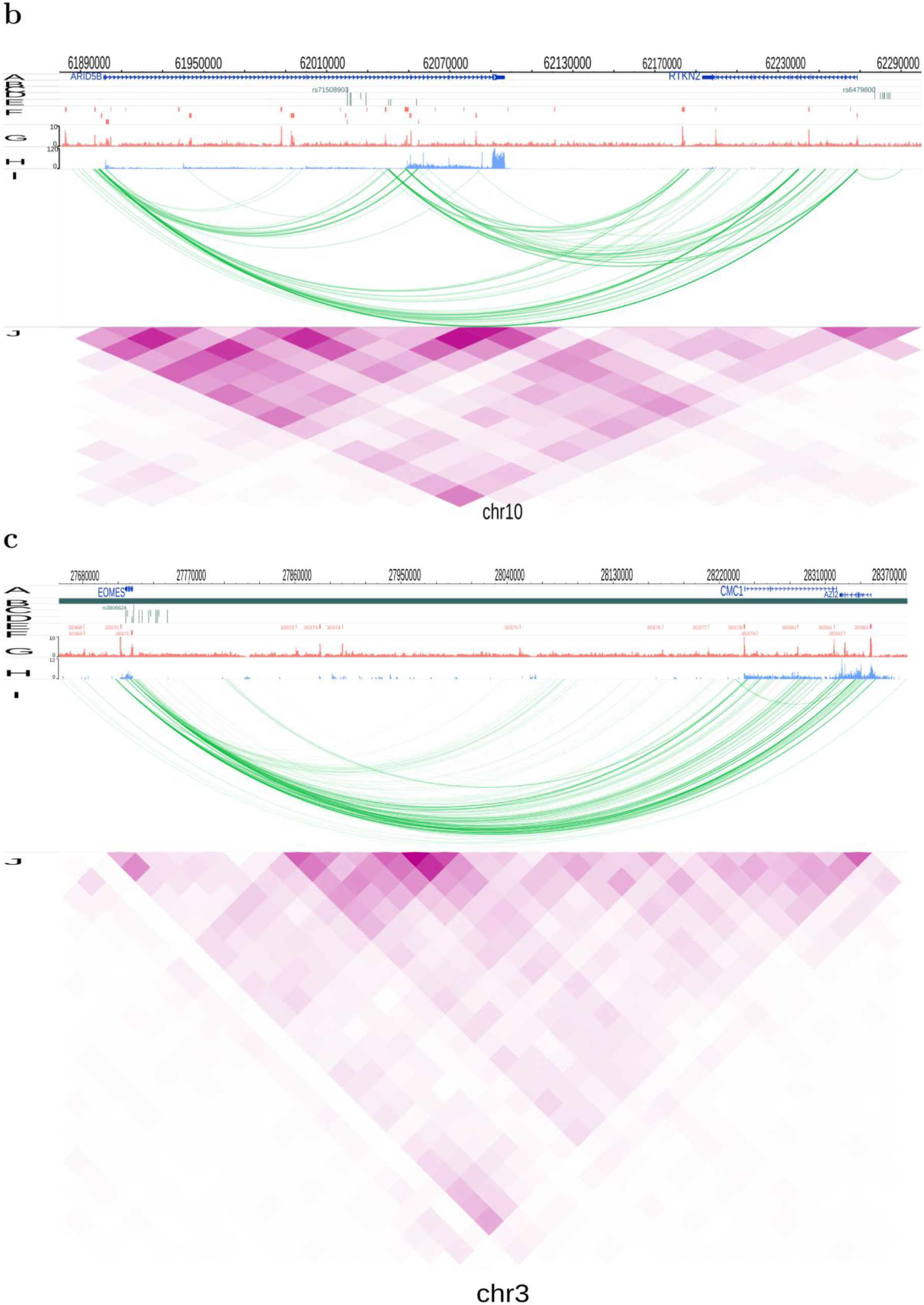
Illustration of genomic interaction activities around three RA associated loci. Screenshots of the SNPs (dark green), ATAC-seq peaks (red), RNA-seq (blue) and CHi-C interactions (green). **a**, RBPJ loci, demonstrating how both interactions between the associated SNPs/gene promoter, and ATAC-seq intensity increase in magnitude over time (0, 20mins, 1hr, 2hr, 24hr, top to bottom) **b**, ARID5B_RTKN2 loci, demonstrating strong interactions between the region intronic of ARID5B and RTKN2. **c**, EOMES_AZI2 loci demonstrating strong interactions between the region intronic of EOMES and AZI2. Yaxis labels used in the picture: A: RefSeq genes, B: TADs, C: Index SNPs, D: LD SNPs, E: 99% credible sets, F:0 min ATAC-seq peaks, G: 0 min ATAC-seq signal, H: 0 min RNA-seq, I: 0 min CHi-C, J: 20 mins ATAC-seq peaks, K: 20 mins ATAC-seq signal, L: 20 mins RNA-seq, M: 20 mins CHi-C, N: 1 hrs ATAC-seq peaks, O: 1 hrs ATAC-seq signal, P: 1 hrs RNA-seq, Q: 1 hrs CHi-C, R: Index SNPs, S: LD SNPs, T: 99% credible sets, U:24 hrs ATAC-seq peaks, V: 24 hrs ATAC-seq signal, W: 24 hrs RNA-seq, X: 24 hrs CHi-C, Y: 24 hrs Hi-C.

### 10. Drug targets

We performed drug target identification of the genes identified by Capture Hi-C as previously described^24^. Briefly, genes showing interactions with RA associated variants were filtered by expression and used to interrogate DrugBank v5.0.11 to find potential drugs. Out of 167 CHi-C genes interacting with RA implicated enhancers, 18 were targets for 49 drugs (similar to our previous findings). Three of these, corresponding to 4 drugs, are already used in RA.

**Supplementary Table 2.** Genes implicated through interactions with RA associated variants in the CD4+ T cell data and associated drugs

| Gene |  | Drug |
| --- | --- | --- |
| AARS2 | DB00160 | L-Alanine |
| CDK6 | DB09073 | Palbociclib |
| CDK6 | DB11730 | Ribociclib |
| CDK6 | DB12001 | Abemaciclib |
| CSF2RB | DB00020 | Sargramostim |
| DGKA | DB00163 | Vitamin E |
| ERBB2 | DB00072 | Trastuzumab |
| ERBB2 | DB01259 | Lapatinib |
| ERBB2 | DB05773 | Trastuzumab emtansine |
| ERBB2 | DB06366 | Pertuzumab |
| ERBB2 | DB08916 | Afatinib |
| ERBB2 | DB12267 | Brigatinib |
| GSN | DB09130 | Copper |
| HDAC2 | DB00227 | Lovastatin |
| HDAC2 | DB00277 | Theophylline |
| HDAC2 | DB00313 | Valproic Acid |
| HDAC2 | DB01223 | Aminophylline |
| HDAC2 | DB01303 | Oxtriphylline |
| HDAC2 | DB02546 | Vorinostat |
| HDAC2 | DB05015 | Belinostat |
| HDAC2 | DB06176 | Romidepsin |
| HDAC2 | DB06603 | Panobinostat |
| HDAC2 | DB09091 | Tixocortol |
| HIF1A | DB01136 | Carvedilol |
| HLA-DQB1 | DB00071 | Insulin Pork |
| HSPA8 | DB01254 | Dasatinib |
| HSPA8 | DB09130 | Copper |
| IL1R1 | DB00026 | Anakinra |
| MAPK14 | DB01254 | Dasatinib |
| MMP9 | DB00143 | Glutathione |
| MMP9 | DB00786 | Marimastat |
| MMP9 | DB01017 | Minocycline |
| MMP9 | DB01197 | Captopril |
| MMP9 | DB01296 | Glucosamine |
| MYC | DB08813 | Nadroparin |
| PIK3CA | DB00201 | Caffeine |
| PIK3CA | DB12483 | Copanlisib |
| PTGDR2 | DB00328 | Indomethacin |
| PTGDR2 | DB00605 | Sulindac |
| PTGDR2 | DB00770 | Alprostadil |
| PTGDR2 | DB00917 | Dinoprostone |
| PTGDR2 | DB01088 | Iloprost |
| SIRT5 | DB04786 | Suramin |
| VEGFA | DB00112 | Bevacizumab |
| VEGFA | DB01017 | Minocycline |
| VEGFA | DB01120 | Gliclazide |
| VEGFA | DB01136 | Carvedilol |
| VEGFA | DB01270 | Ranibizumab |
| VEGFA | DB03754 | Tromethamine |
| VEGFA | DB05294 | Vandetanib |
| VEGFA | DB06779 | Dalteparin |
| VEGFA | DB08885 | Aflibercept |
| VEGFA | DB09301 | Chondroitin sulfate |

### 11. CRISPR experiments in HEK293T cells

HEK293T cells were selected as a suitable model system as they displayed a similar genetic architecture, in terms of TADS across the region, to our primary T cell data (Supplementary Fig. 14 a). CRISPR guides were designed to cover the single enhancer that contains the 2 ATAC-seq peaks and 3 SNPs associated with RA, pooling the guides in a single transfection (Supplementary Fig. 14 b).

**Supplementary Fig. 14.**
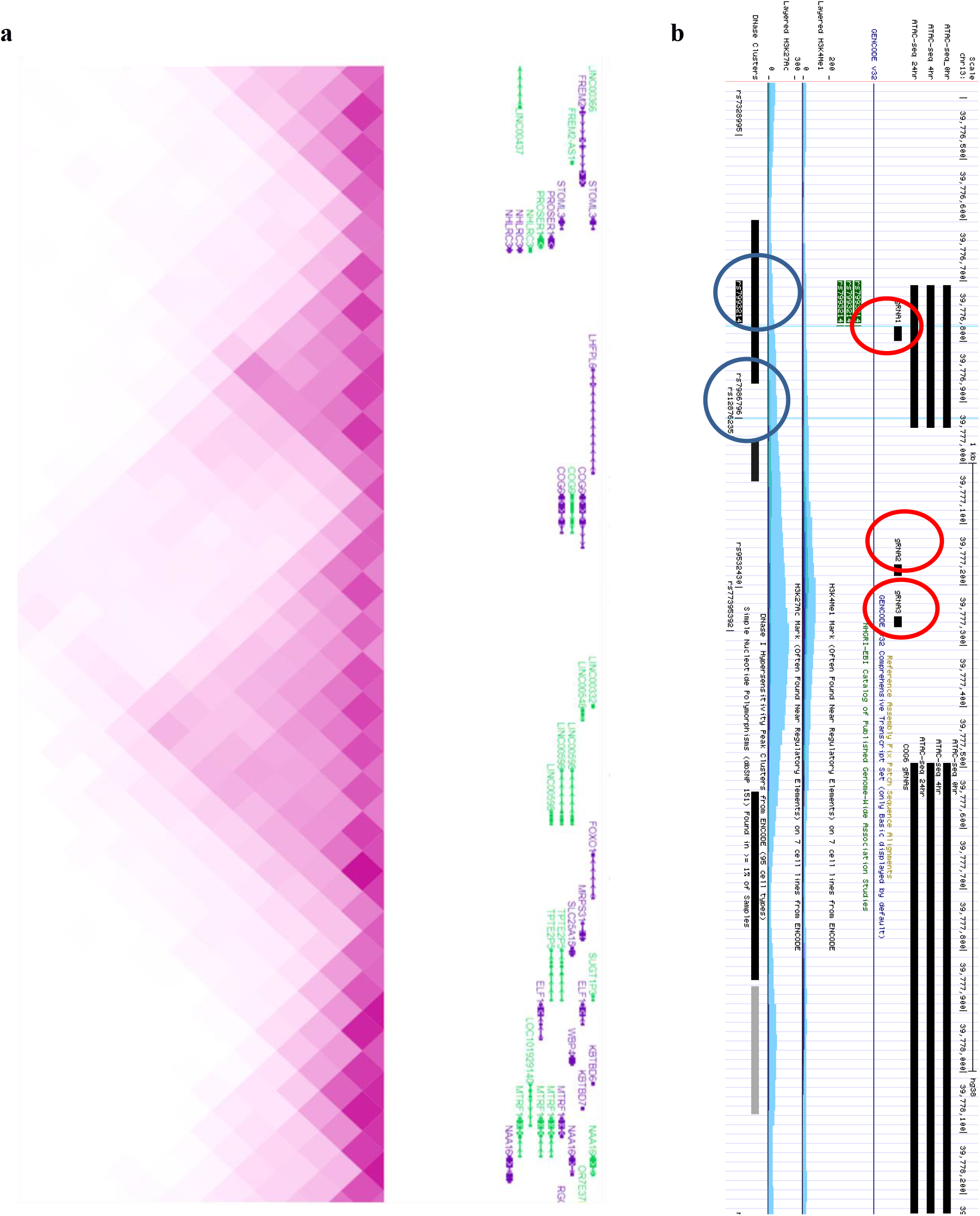
Demonstrating the TAD structure across the COG6/FOXO1 region (a) and the positioning of the gRNA CRISPR guides (red circles) in relationship to the associated SNPs (blue circles), ATAC-seq peaks and enhancer region (b).

Supplementary Table 3 **ALL Autoimmune loci.** Table of all ATAC-seq peaks containing a SNP in the 99% credible set for an autoimmune disease, the promoters they interact with and the correlation between ATAC-seq activity, interaction strength and gene expression. Included as excel spreadsheet.

Supplementary Table 4 **ALL RA loci.** Table of all ATAC-seq peaks containing a SNP in the 99% credible set for RA, the promoters they interact with and the correlation between ATAC-seq activity, interaction strength and gene expression, plus SNPs with eQTL evidence. Included as excel spreadsheet.

